# IL-6/STAT3 signaling drives early-stage pancreatic cancer cachexia via suppressed ketogenesis

**DOI:** 10.1101/2022.10.18.512778

**Authors:** Paige C. Arneson-Wissink, Heike Mendez, Katherine Pelz, Jessica Dickie, Alexandra Q. Bartlett, Beth L. Worley, Stephanie M. Krasnow, Robert Eil, Aaron J. Grossberg

## Abstract

Cancer cachexia is highly prevalent in patients with pancreatic ductal adenocarcinoma (PDAC). Although advanced cachexia is associated with inflammatory signaling, the early events driving wasting are poorly defined. Using an orthotopic mouse model of PDAC, we find that early cachexia is defined by a pronounced vulnerability to undernutrition, characterized by increased skeletal muscle wasting. PDAC suppresses lipid beta oxidation and impairs ketogenesis in the liver, which coordinates the adaptive response to nutritional scarcity. When PDAC mice are fed ketogenic diet, this effect is reversed, and muscle mass is preserved. Furthermore, physiologic levels of ketones are sufficient to protect myotubes against PDAC-associated wasting. Interleukin-6 (IL-6) drives liver metabolic reprogramming, and hepatocyte-specific loss of Signal Transducer and Activator of Transcription 3 (STAT3) is sufficient to prevent PDAC-associated muscle loss. Together, these studies define a key role for the liver in cachexia development and directly link skeletal muscle homeostasis to hepatic lipid oxidation.

## INTRODUCTION

Cachexia is a widely-documented co-morbidity of cancer that negatively impacts patient quality of life and their ability to receive life-extending anti-cancer therapies, and yet there are no efficacious treatments for cachexia (Fearon et al., 2013). Patients with pancreatic ductal adenocarcinoma (PDAC) are especially at risk; about 80% of PDAC patients will develop cachexia at some point during their cancer journey (von Haehling et al., 2016). Cancer cachexia is a complex behavioral and metabolic syndrome hallmarked by accelerated fat and muscle wasting, often in the context of nutritional deficit (Baracos et al., 2018; Olson et al., 2021). Cachexia differs from pure undernutrition in two key ways: 1) skeletal muscle is relatively spared during periods of nutritional scarcity, but not in cachexia(Cahill, 1970); 2) nutritional supplementation is inadequate to reverse cachexia (Omlin et al., 2013). Furthermore, cachectic mice present with metabolomic and transcriptomic signatures that differ from fasted mice, outlining unique mechanisms of tissue catabolism between starvation and cachexia (Goncalves et al., 2018). Given this fundamental difference between undernutrition and cachexia, significant effort has been directed to understanding the paraneoplastic processes underlying the persistent and pervasive wasting associated with cachexia. Features common across multiple models of both cancer and non-cancer cachexia include increased resting metabolic rate, elevated lipolysis, and enhanced skeletal muscle proteolysis (Fearon et al., 2012; Lieffers et al., 2009; Penna et al., 2019; Pettersen et al., 2017).

Decades of research defined the unique and powerful role of inflammation as a regulator of acute sickness responses and driver of cachexia in patients and rodents (Baracos *et al*., 2018; Fearon *et al*., 2013; Fearon *et al*., 2012; Fearon et al., 2006; Grossberg et al., 2010). Administration of inflammatory cytokines is sufficient to induce the behavioral and metabolic changes characteristic of cachexia, and blockade of cytokine signaling prevents the onset of experimental cachexia in several models (Acharyya et al., 2004; Braun et al., 2013; Braun et al., 2012; Fujita et al., 1996; Llovera et al., 1998; Zhu et al., 2018). Cytokines act on the brain to suppress food intake and increase thermogenesis, adipose tissue to promote “browning” and lipolysis, and muscle to promote proteolysis (Emmerson et al., 2017; Grossberg *et al*., 2010; Mullican et al., 2017; Scarlett et al., 2007). Of the myriad of cytokines implicated in the development of cachexia, interleukin (IL)-6, specifically, is of particular interest in PDAC, as it is associated with disease progression, metastasis, mortality, and cachexia (Babic et al., 2018; Ebrahimi et al., 2004; Martignoni et al., 2005; Okada et al., 1998; Ramsey et al., 2019). Elevated IL-6 levels are associated with weight loss in murine PDAC, and IL-6 drives cachexia in murine models of PDAC and other cancers (Baltgalvis et al., 2008; Bonetto et al., 2012; Bonetto et al., 2011; Flint et al., 2016; Narsale and Carson, 2014; Rupert et al., 2021; Suh et al., 2013). Overexpression of IL-6 elicits skeletal muscle wasting *in vivo*, and IL-6 induces lipolysis and wasting in adipocytes *in vitro* (De Benedetti et al., 2002; Ji et al., 2011; Trujillo et al., 2004). However, to date anti-cytokine approaches have failed to yield clinically efficacious treatment strategies in patients. This may be due to a focus on late-stage cachexia, which is often associated with advanced cancer and multiple comorbidities (Fearon et al., 2011).

Cachexia is clinically defined by certain thresholds of weight and muscle loss (Fearon *et al*., 2011), and most studies evaluating cancer cachexia compare mice with fully developed cachexia against healthy control populations (Arneson-Wissink et al., 2020; Langer et al., 2022; Michaelis et al., 2017; Talbert et al., 2019). Because of the focus on advanced disease, the pre-cachectic state remains poorly defined. The early biology driving initial wasting is not well-characterized and may differ from the inflammation-driven biology defined in more advanced cachexia (Dalise et al., 2020; Fearon *et al*., 2011; Zwickl et al., 2020). Indeed, the link between inflammation and cachexia is largely based on elevations in circulating cytokines observed when significant wasting has already occurred (Bonetto *et al*., 2012; Flint *et al*., 2016; Onesti and Guttridge, 2014). A recent study of early murine PDAC cachexia, using the autochthonous KPC model, revealed no elevations in cachexia-associated cytokines, and was unable to discriminate cachexia from pure undernutrition (Danai et al., 2018). Accordingly, visceral adipose tissue from patients with PDAC cachexia does not exhibit gene expression consistent with browning or enhanced lipolysis (Xu et al., 2022). These data indicate that the processes canonically associated with cachexia may not be the driving processes most relevant during the pre-cachectic state.

Maintaining metabolic homeostasis requires rapid adaptations to variable nutrient conditions. The liver is central to these adaptations by storing excess nutrition (fat and glycogen) during periods of nutritional abundance and distributing resources during nutritional scarcity. In healthy animals, during times of low nutritional availability, the liver shifts metabolic activity from glucose storage to gluconeogenesis and ketogenesis (Felig et al., 1969; Puchalska and Crawford, 2017; Rothman et al., 1991). This results in a supply of energy-rich molecules (glucose and ketone bodies, respectively) that are utilized by the brain, heart, and skeletal muscle. The oxidation of lipids by the liver is thought to be central to skeletal muscle preservation during fasting (Cahill, 1970). This hypothesis is supported by the observations that exhaustion of adipose stores precipitates rapid muscle catabolism in humans and rodents undergoing prolonged starvation (Flatt and Quail, 1981; Goodman et al., 1980; Goodman et al., 1984; Goodman and Ruderman, 1980). Pharmacologic inhibition of lipolysis or fatty acid oxidation (Lowell and Goodman, 1987), or damage to the liver with the hepatotoxin CCl_4_ (Flatt and Quail, 1981) each independently accelerated muscle wasting in fasted rats, together implicating hepatic lipid mobilization and oxidation in skeletal muscle conservation. Furthermore, hepatocytes express cytokine receptors, which allows the liver to coordinate a systemic response to inflammation by initiating the acute phase protein response to inflammatory signals. Existing work shows that cancer associates with impaired ketogenesis, and that restored ketone production ameliorates muscle wasting in other instances of cancer cachexia (Flint *et al*., 2016; Goncalves *et al*., 2018; Shukla et al., 2014). Though the liver is well-positioned to coordinate altered metabolism in cachexia, its role remains poorly defined, particularly early in cancer cachexia, when wasting is initiated.

We propose that PDAC causes hepatic metabolic changes that underlie wasting during cachexia development. We leverage an orthotopic PDAC (OT-PDAC) mouse model of early cachexia to define the events that mediate early-stage tissue wasting, and examine the roles of under-nutrition, hepatic adaptive metabolism to undernutrition, and inflammatory signaling in driving PDAC cachexia. Using this approach, we found that disrupted hepatic lipid metabolism is a key feature of early-stage cachexia, and that reversal of this deficit using two genetic models that block IL-6 signaling, or a ketogenic diet intervention, were sufficient to prevent PDAC-associated muscle loss.

## RESULTS

### PDAC elicits a metabolic vulnerability to nutritional stress

We modeled PDAC cachexia in mice using orthotopic implantation of PDAC cancer cells derived from the *Kras*^*G12D/+*^; *Tp53*^*R172H/+*^; *Pdx-cre* (KPC) mouse into the pancreatic tail (Michaelis *et al*., 2017). A*d libitum* fed PDAC-bearing mice experience progressive wasting over the first two weeks of tumor development, which paralleled a decrease in food intake (Figure S1A-E). This led us to question whether tissue wasting was solely attributable to undernutrition or malabsorption, as previously suggested (Danai *et al*., 2018). If PDAC cachexia is driven by a paraneoplastic process, mice would have increased vulnerability to nutritional stress, such as food restriction. We tested this hypothesis by challenging mice with 50% food-restriction (FR) for 4 days prior to termination on day 7, before we see evidence of wasting among *ad libitum* fed mice (Figure 1A). Diet availability did not affect the terminal size of pancreatic tumors (Figure 1B). Although no differences in fat mass were observed between sham and PDAC mice maintained on either diet, FR elicited a loss of both fat free mass (measured by echoMRI) and skeletal muscle mass in PDAC, but not sham, mice (Figure 1C-D). Gross tissue weights of gonadal white adipose tissue (gWAT) and gastrocnemius muscle also showed a biased loss in muscle only in FR PDAC mice (Figure 1E-F). Similarly, expression of atrophy-associated genes *Fbxo32, Foxo1*, and *Trim63* increased in gastrocnemius muscle from FR PDAC mice, but not in the *ad libitum* PDAC mice (Figure 1G). The application of an alternative nutritional challenge, an acute 24-hour fast, revealed a similar elevation in *Fbxo32, Foxo1*, and *Trim63* expression in PDAC compared to sham mice (Figure S1F). Together, these data show that PDAC mice exhibit an increased vulnerability to nutritional stress, characterized by lean muscle wasting.

**Figure 1.**
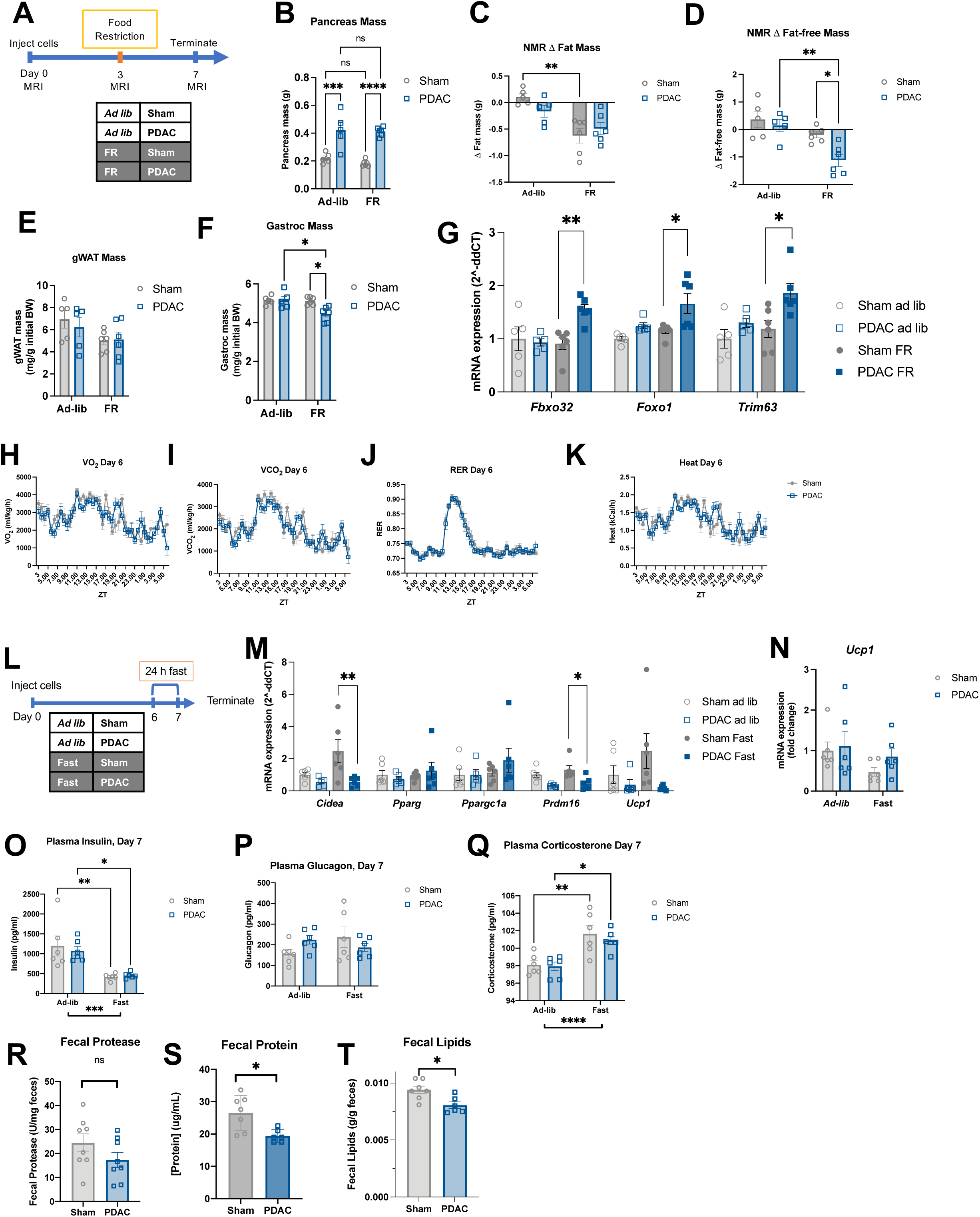
PDAC elicits a metabolic vulnerability to nutritional stress. (A) Schema of food restriction (FR) pre-cachexia model used in panels B-G. n = 3 female, 2 male (sham *ad libitum*); 2 female, 4 male (sham FR); 3 female, 2 male (PDAC *ad libitum*); 2 female, 4 male (PDAC FR). (B) Pancreas mass at termination of PDAC and sham animals. Change in fat mass (C) and fat-free mass (D) relative to baseline at 7 d post implantation, measured by MRI. gWAT (E) and gastrocnemius (F) mass at termination of PDAC and sham animals. (G) qPCR analysis of muscle atrophy genes (*Foxo1, Fbxo32, Trim63*) measured in gastrocnemius muscle. (H-K) Indirect calorimetry during d 6 post-implantation, representing final 24 hours before termination timepoint. n = 4 male and 4 female mice per group. (H) VO_2_, (I) VCO_2_, (J) respiratory exchange ratio (RER), (K) heat output. (L) Schema of the 24 h fast pre-cachexia model used in panels M-T. n = 3 female, 3 male mice per group (M-Q); n = 4 female, 4 male mice per group (R); n = 3 female, 4 male (sham), 3 female, 3 male (PDAC) (S-T). (M) qPCR analysis of adipose browning genes (*Cidea, Pparg, Ppargc1a, Prdm16, Ucp1*) measured in white adipose tissue. (N) *Ucp1* expression in brown adipose tissue (BAT). (O) Terminal plasma insulin, (P) glucagon, and (Q) corticosterone measured in *ad libitum* and fasted sham and PDAC mice. (R) Fecal protease activity, (S) fecal protein content, and (T) fecal lipid content, normalized to mass of feces of sham and PDAC animals fed *ad libitum* for 7 days. Error bars represent SEM. All analyses for 2×2 studies were statistically tested with a 2-way ANOVA with Tukey correction for multiple comparisons. Indirect calorimetry measures were statistically tested with repeated measures ANOVA with Šidák correction for multiple comparisons. Pairwise comparisons statistically tested with parametric, unpaired *t* test. **** p<0.0001, ***p<0.001, **p<0.01, *p<0.05. Independent biological replicates are indicated by individual points on bar graphs, unless noted otherwise.

To identify the source of this metabolic vulnerability, we first examined the impact of PDAC on basal metabolism (Falconer et al., 1994; Lieffers *et al*., 2009). Indirect calorimetry revealed no differences in VO2, VCO2, respiratory exchange ratio (RER), or heat generation between PDAC and pair-fed sham mice at the same timepoint (Figure 1H-K). We neither observed elevated expression of browning-associated genes (*Cidea, Pparg, Ppargc1a, Prdm16, Ucp1*) in gWAT, nor the mitochondrial uncoupler *Ucp1* in brown adipose tissue (BAT) at 7-or 10-days post implantation (Figure 1L-N, S1G). Plasma insulin, glucagon, and corticosterone were also not perturbed in PDAC mice, suggesting tissue wasting is not mediated by endocrine dysregulation (Figure 1O-Q). Finally, we did not find a difference in fecal protein, lipid, or protease levels at 7 days post implantation, in agreement with previously published data in the same model from 10 days post implantation (Figure 1R-T)(Olson *et al*., 2021). This series of data show that sensitivity of PDAC mice to nutritional stress is not driven by BAT thermogenesis, adipose browning, endocrine dysfunction, or malabsorption.

### PDAC dysregulates hepatic metabolic adaptation to undernutrition

The liver orchestrates the adaptive response to undernutrition by increasing lipid utilization and releasing ketone bodies as a fuel source to the brain, heart, and muscle. It is postulated that lipid oxidation by the liver is essential for lean tissue sparing during fasting, as evidenced by the dependence of muscle preservation on adequate lipid stores and normal hepatic function(Cahill, 1970; Parrilla, 1978). Thus, we examined hepatic metabolism in fasted, early-stage PDAC mice (Figure 2A). First, we observed that PDAC mice had lower fasting blood ketone levels than sham mice, and that there was not a difference in ketone body redox state between sham and PDAC mice (Figure 2B, S2A-C). We then challenged fasted PDAC and sham mice with a bolus of the medium-chain fatty acid octanoate, which revealed impaired ketogenic capacity in PDAC mice (Figure 2C-D). We next assessed liver metabolism on a molecular level by measuring the expression of key enzymes involved in liver lipid import (*Cpt1)*, beta oxidation (*Acadm, Acadl, Acat1, Acox1, Acsl1, Echs1, Ehhadh)*, ketogenesis (*Acaa2, Bdh1, Hmgcl, Hmgcs2*) and the master regulator of hepatic lipid metabolism, *Ppara*. We saw downregulation of all of these transcripts in PDAC mice compared to sham mice, except *Acat1* (Figure 2E-F). Although the expression of lipid oxidation genes can be induced by lipid availability, we observed no differences in plasma levels of triglycerides or non-esterified fatty acids (NEFAs) at either 7-or 10-days post implantation (Figure S2D-G). Furthermore, we did not see differences in lipid content in gastrocnemius muscle or whole liver tissue, indicating that the decrease in beta oxidation did not yield lipid buildup in either tissue (Figure S2H-P). To evaluate whether decreased ketogenesis was solely an artifact of our orthotopic PDAC model, we looked at fasting plasma metabolites in the autochthonous PDAC, *Kras*^*G12D/+*^; *Tp53*^*R172H/+*^; *Pdx-cre* (KPC) mouse model of PDAC as well as in samples from patients with localized, resected PDAC collected at the time of surgery, when they were instructed to fast for at least 8 hours. Ketogenesis was suppressed in all patients with PDAC, compared with patients with non-malignant indications for surgery, as well as in KPC mice (Figure 2G-H, Table S1). These data show that impaired hepatic lipid metabolism, and ketogenesis, in particular, is pervasive across both mouse models of PDAC and human patient populations.

**Figure 2.**
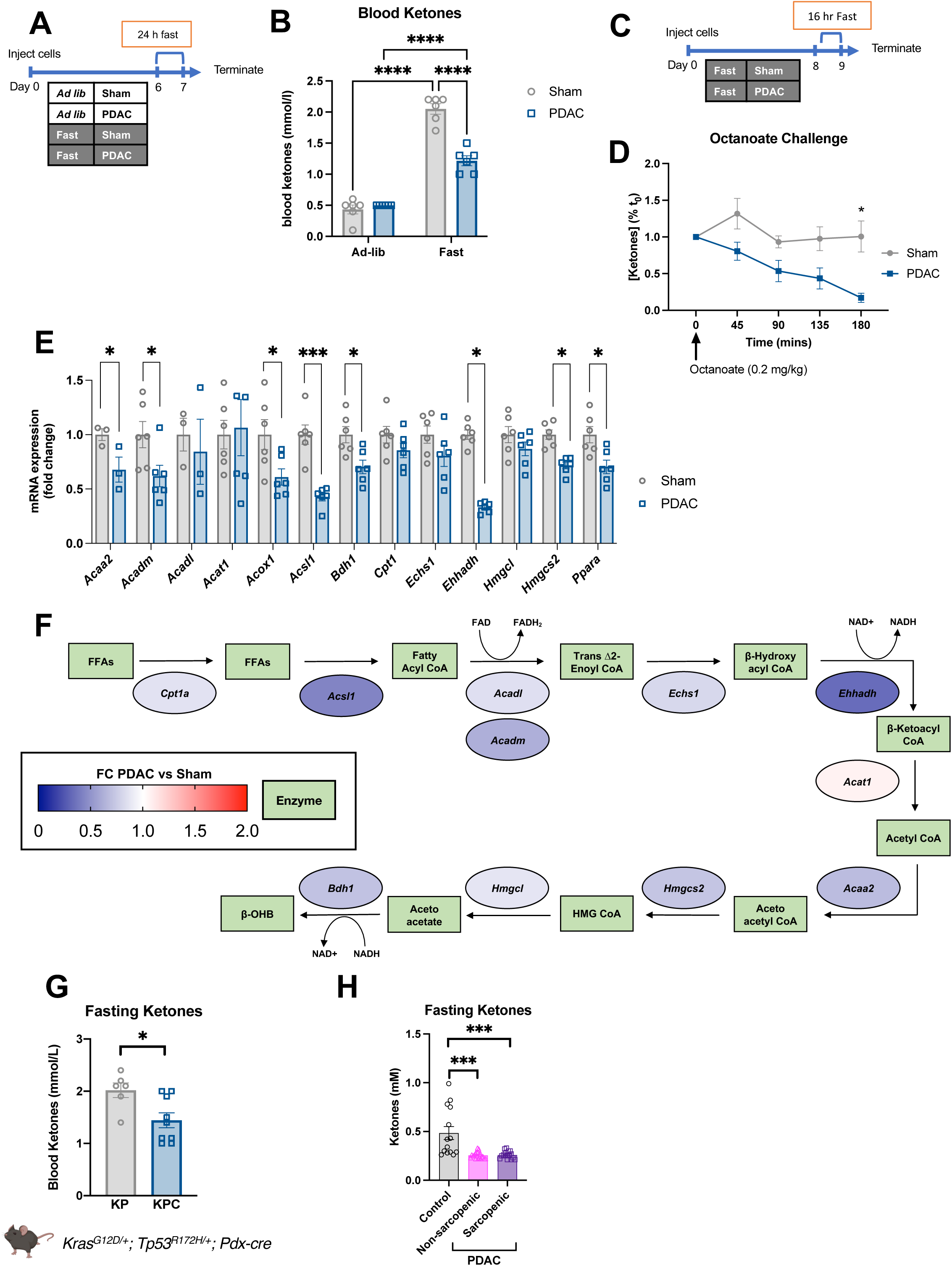
PDAC dysregulates hepatic metabolic adaptation to undernutrition. (A) Schema of pre-cachexia model with 24 h fast used in panels B, E-F. n = 3 female, 3 male mice per group. (B) Blood ketone levels in *ad libitum* and fasted sham and PDAC mice. (C) Schema for octanoate challenge after 16 h fast, used in panel D. n = 4 female, 4 male (PDAC); 3 female, 4 male (sham). (D) Octanoate challenge blood ketone levels in sham and PDAC mice. n = 7 sham, 8 PDAC. (E) qPCR analysis of lipid metabolism genes (*Acaa2, Acadm, Acadl, Acat1, Acox1, Acsl1, Bdh1, Cpt1, Echs1, Ehhadh, Hmgcl, Hmgcs2, Ppara*) measured in liver tissue. (F) Schematic depicting the relative expression of hepatic beta oxidation and ketogenic genes from panel D, with color representing log_2_ fold-change (PDAC v sham). (G) Fasting ketones measured in autochthonous KPC mice and littermate KP controls, collected from 12-13 week old mice after 24 h fast. n = 3 female, 3 male (KP); 7 female, 2 male (KPC). (H) Fasting ketones measured in plasma from non-sarcopenic and sarcopenic patients with PDAC at time of laparotomy. n = 7 female control, 13 female PDAC no sarcopenia, 10 female PDAC sarcopenia, 7 male control, 7 male PDAC no sarcopenia, 8 male PDAC sarcopenia. Controls in (H) collected from patients undergoing endoscopic ultrasound for non-malignant indications. Error bars represent SEM. All analyses for 2×2 studies were statistically tested with 2-way ANOVA with Tukey correction for multiple comparisons. Octanoate challenge was statistically tested with repeated measures ANOVA with Šidák correction for multiple comparisons. Pairwise comparisons tested with parametric, unpaired *t* test. 3-group analyses tested with one-way ANOVA and Tukey correction for multiple comparisons. **** p<0.0001, ***p<0.001, **p<0.01, *p<0.05. Independent biological replicates are indicated by individual points on bar graphs, unless noted otherwise.

### Ketogenic diet prevents muscle and liver dysfunction in PDAC cachexia

Our observations that circulating ketones were lower across several PDAC data sets led us to question whether lower ketone levels were detrimental to muscle, and if restored ketogenesis would be protective against cachexia. Although PDAC mice exhibit a deficit in fasting ketogenesis, we questioned whether chronic exposure to a ketogenic diet would yield sufficient ketones to preserve muscle. To test this hypothesis, we fed PDAC and sham mice ketogenic diet or calorically matched control diet for 10 days following implantation (Tables S2-3). We pair-fed sham mice to PDAC given the same diet to focus our study on tumor-driven, rather than calorie-driven effects on muscle and liver (Figure 3A). Ketogenic diet did not affect the size of tumor, as measured by gross pancreas mass at termination (Figure 3B). We did not see differences in cumulative caloric intake, body mass, gross liver mass, or gross gWAT mass between any of the groups (Figure 3C-D, Figure S3A-F). Ketogenic diet induced elevated blood ketone levels in all mice and slight increases in plasma triglyceride levels (Figure 3E, S3G). As expected, all ketogenic diet-fed mice had lowered blood glucose levels, and PDAC mice in both groups had slightly lower blood glucose than sham animals (Figure 3F). Ketogenic diet reversed PDAC-associated muscle wasting, measured as both gastrocnemius and tibialis anterior (TA) gross muscle mass and TA myofiber cross-sectional area (Figure 3G-H, J, S3H-J). Paradoxically, ketogenic diet did not reverse the increased expression of atrophy markers: *Trim63, Foxo1*, and *Fbxo32* (Figure 3I). Because blood ketone levels were restored in ketogenic-diet fed PDAC mice, we next looked at the expression of hepatic lipid metabolism genes. We found that ketogenic diet restored the expression of several genes that were downregulated in control diet PDAC mice, including *Acadl*, a dehydrogenase essential for fatty acid oxidation, *Acaa2*, the final step in mitochondrial beta oxidation, *Hmgcs2*, which catalyzes the rate limiting step in ketogenesis, and *Bdh1*, which catalyzes the interconversion between the ketone bodies acetoacetate and beta hydroxybutyrate (Figure 3K). The restoration of ketogenesis and gene expression indicate that the removal of carbohydrates from the diet is sufficient to reverse PDAC-induced metabolic reprogramming. Whether ketogenesis, and ketones, specifically, mediated the preservation of skeletal muscle remained unaddressed by this study.

**Figure 3.**
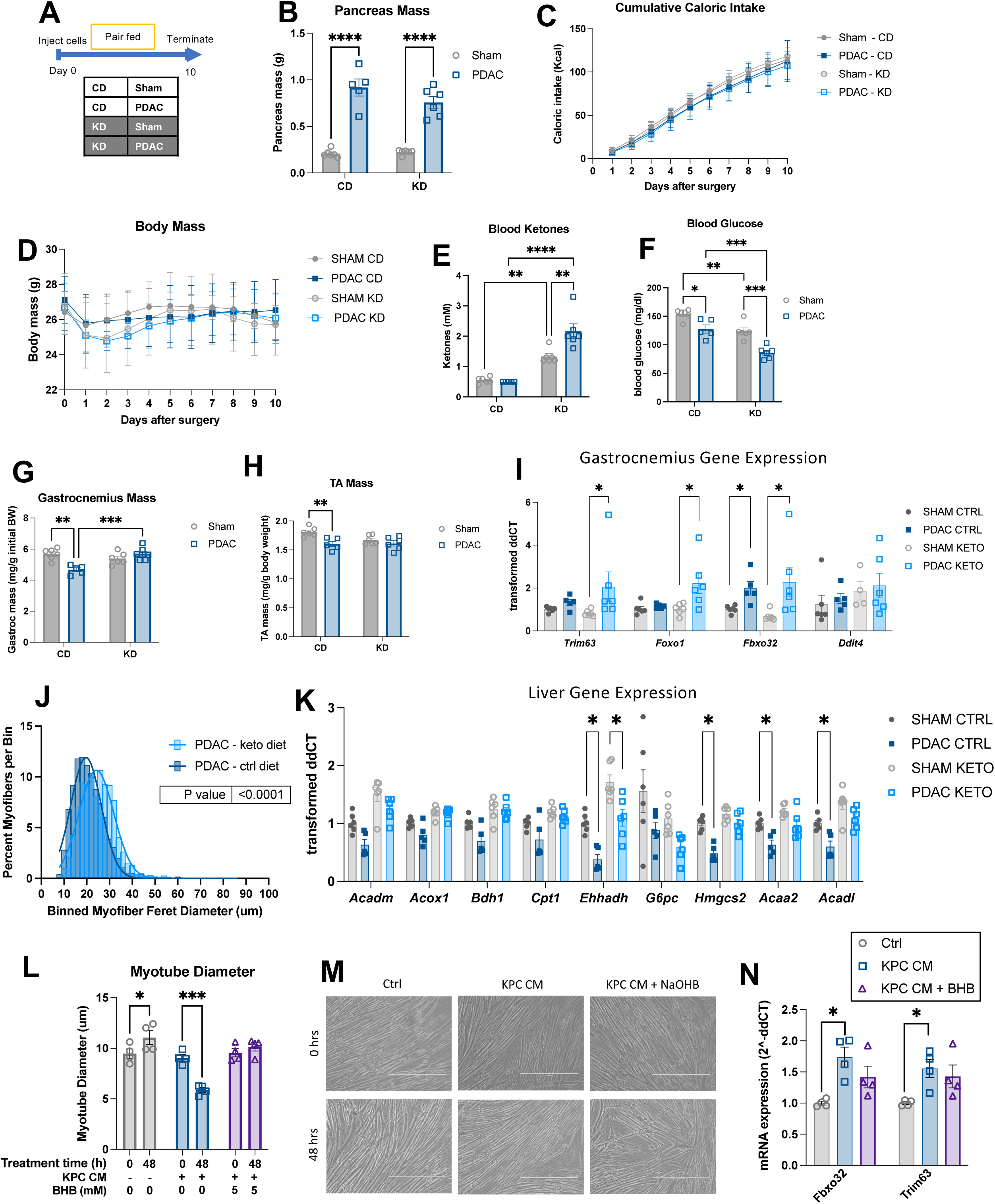
Ketones reverse myotube atrophy *in vitro* and muscle wasting in PDAC mice. (A) Timeline schematic for *in vivo* ketogenic diet study used in panels G-P. CD = control diet, KD = ketogenic diet. n = 6 male mice (sham/CD, sham/KD, PDAC/KD), 5 male mice (PDAC/CD). (B) Pancreas mass at termination of PDAC and sham animals. (C) Cumulative caloric intake over study. (D) Body mass throughout study. Blood ketone (E) and blood glucose (F) levels in CD and KD, sham and PDAC animals at termination. (G) Gastrocnemius muscle mass normalized to initial body weight. (H) Terminal tibialis anterior muscle mass normalized to initial body weight. (I) qPCR analysis of muscle atrophy genes (*Foxo1, Fbxo32, Trim63, Ddit4*) measured in gastrocnemius muscle. (J) Quantification of minimum feret diameter of myofibers from PDAC mice fed control or ketogenic diet. Feret diameters were binned to a histogram and fit with a non-linear regression (Gaussian, least squares regression). Myofibers in ketogenic diet PDAC mice were significantly larger; P < 0.0001 by extra sum-of-squares F test. (K) qPCR analysis of lipid metabolism genes (*Acaa2, Acadl, Acadm, Acox1, Bdh1, Cpt1, Ehhadh, G6pc, Hmgcs2*) measured in liver tissue. (L) C2C12 myotube diameter at baseline and after 48 h exposure to KPC conditioned media (KPC CM) ± sodium beta hydroxybutyrate (BHB). n = 4 wells from 6-well plates per group. (M) Phase contrast images of C2C12 myotubes at 0 and 48 hours in indicated media treatments. Scale bar = 400 μm. (N) qPCR analysis of E3 ubiquitin ligases *Trim63* and *Fbxo32* in C2C12 myotubes 48 hours after media change. Error bars represent SEM. All analyses for 2×2 studies were statistically tested with 2-way ANOVA with Tukey correction for multiple comparisons. 3 and 4-group analyses tested with one-way ANOVA and Tukey correction for multiple comparisons. **** p<0.0001, ***p<0.001, **p<0.01, *p<0.05. Independent biological replicates are indicated by individual points on bar graphs, unless noted otherwise.

We tested this hypothesis by assessing the impacts of sodium beta-hydroxybutyrate (BHB) on atrophic C2C12 myotubes. BHB is the predominant ketone body produced during hepatic ketogenesis. To assess the capacity of BHB to reverse muscle wasting, we developed an *in vitro* model of PDAC-associated myotube atrophy by exposing differentiated C2C12 myotubes to KPC cell conditioned media (KPC CM). KPC CM-treated myotubes showed a significant decrease in diameter after 24 h of treatment, which was abrogated in a dose dependent manner with increasing concentrations of BHB (Figure S3K). After either 24-or 48-hour treatment, a physiologic concentration (5 mM) of BHB was sufficient to prevent myotube atrophy induced by KPC CM (Figure 3L-M, S3L). While KPC CM treatment increases myotube expression of E3 ubiquitin ligases *Trim63* and *Fbxo32*, as we saw in mice fed ketogenic diet, increased expression of atrophy-associated genes was not reversed with BHB treatment (Figure 3N, S3M). These data indicate that ketone bodies are sufficient to prevent myotube atrophy independent of elevated E3 ubiquitin ligase expression.

### IL-6 expression in the tumor microenvironment drives PDAC cachexia

We were next interested in understanding the nature of the signal to the liver that drives metabolic reprogramming. Interleukin 6 (IL-6) has been identified as a driver of PDAC cachexia in numerous studies and prior work suggests that IL-6 is able to decrease hepatic ketogenesis via signaling through IL-6R (Baltgalvis *et al*., 2008; Bonetto *et al*., 2012; Flint *et al*., 2016; Narsale and Carson, 2014). We observed increased plasma IL-6 in our OT-PDAC model and in PDAC patients (Figure 4A-B). Thus, we sought to elucidate the impacts of IL-6 on hepatic metabolism and ketogenesis. The KPC cells we used in OT-PDAC did not release large amounts of IL-6, and when we implanted these cells into IL-6^−/-^ mice, we did not detect IL-6 in the pancreas (Figure 4C-D). Therefore, we next asked if whole-body IL-6 knock out (IL-6^−/-^) mice would be refractory to cachexia development and ketogenic impairment. To test this, we implanted wild type (WT) and IL-6^−/-^ mice with PDAC tumors and maintained them on *ad libitum* diet for 11 days, at which timepoint WT mice have fully developed cachexia (Figure 4E). We did not measure any significant differences in food intake, body mass, tumor size, or gWAT mass across genotypes (Figure 4F-I). However, IL-6^−/-^ mice were resistant to PDAC-induced muscle wasting, as evidenced by preservation of gastrocnemius muscle mass (Figure 4J). This was supported by molecular data, which showed that elevated expression of muscle catabolism genes (*Fbxo32, Foxo1, Trim63*) was abrogated in the IL-6^−/-^ mice, as compared to WT PDAC mice (Figure 4K). Together these data demonstrate that IL-6 is necessary for the development of cachexia in our PDAC model and broadly support prior research showing that IL-6 is a major contributor to PDAC cachexia.

**Figure 4.**
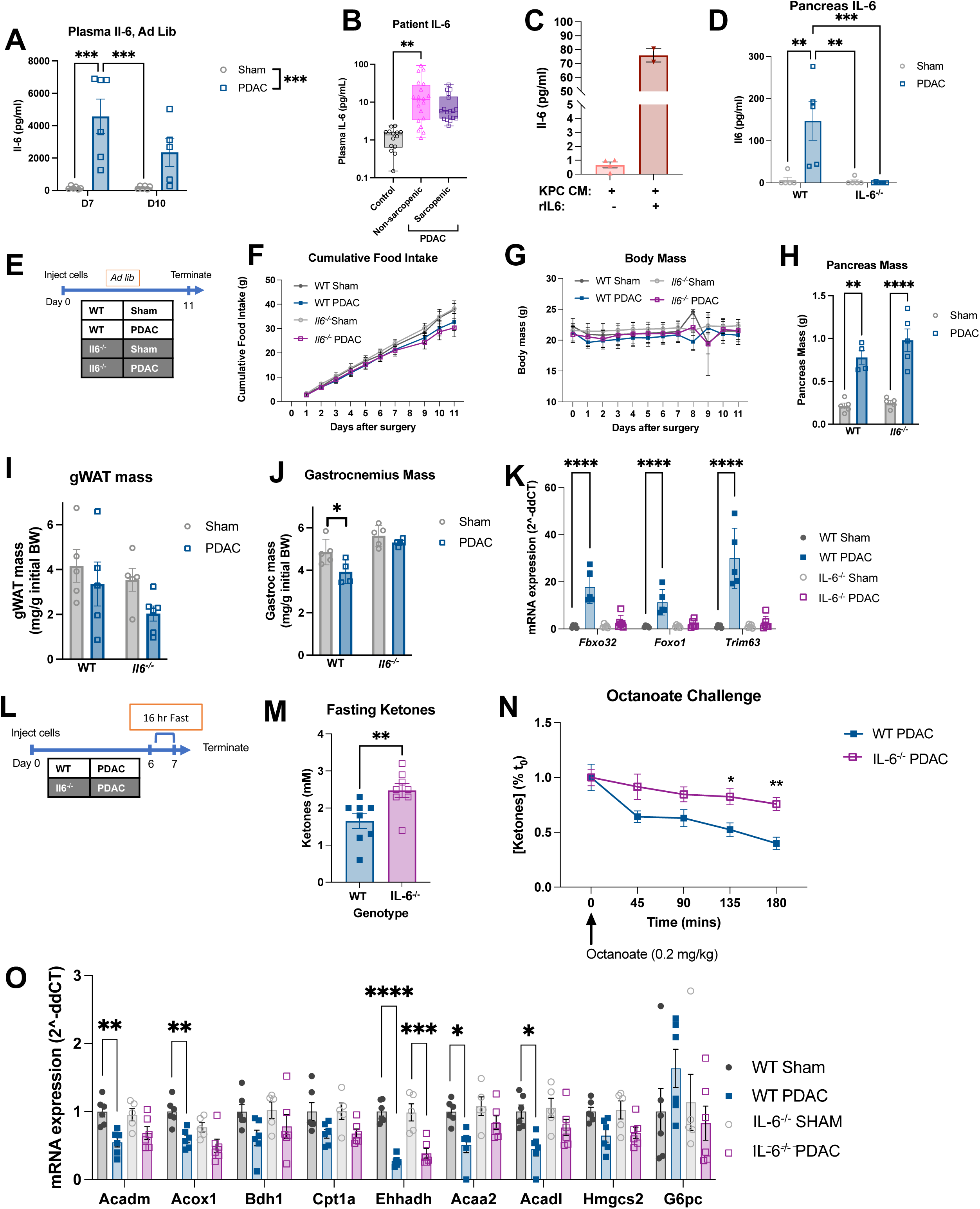
IL-6 expression in the tumor microenvironment drives PDAC cachexia. (A) Plasma IL-6 at 7 d and 10 d post PDAC implantation in mice fed *ad libitum*. n = 3 female, 3 male mice (sham D7, PDAC D7, sham D10); 2 female, 3 male mice (PDAC D10). (B) IL-6 protein measured in plasma from control, non-sarcopenic and sarcopenic patients with PDAC at time of laparotomy. n = 7 female control, 13 female PDAC no sarcopenia, 10 female PDAC sarcopenia, 7 male control, 7 male PDAC no sarcopenia, 8 male PDAC sarcopenia. Controls in (B) collected from patients undergoing endoscopic ultrasound for non-malignant indications. (C) IL6 concentration in KPC CM samples with and without recombinant IL6 spiked in. n = 4 independent media collections (-rIL-6) and 2 independent media collections (+rIL6). (D) IL-6 concentration in whole protein lysate from pancreas of sham and PDAC in wild type and IL-6^−/-^ mice. (E) Timeline schematic outlining study shown in panels D-K. n = 2 female, 3 male (sham/WT); 2 female, 3 male (sham/IL-6^−/-^); 3 female, 2 male (PDAC/WT); 3 female, 3 male (PDAC/IL-6^−/-^). (F) Cumulative food intake, (G) and body mass over study. (H) Pancreas mass at termination of PDAC and sham animals. (I) gWAT and (J) gastrocnemius mass at termination, normalized to initial body weight. (K) qPCR analysis of muscle atrophy genes (*Foxo1, Fbxo32, Trim63*) measured in gastrocnemius muscle. (L) Timeline schematic showing the 16-hour fast paradigm used in panels L-N. n = 8 male mice per group. (M) Blood ketone levels (mM) at termination. (N) Blood ketone levels in WT and IL-6^−/-^ PDAC mice in response to octanoate challenge after 16 h fast. (O) qPCR analysis of lipid metabolism genes (*Acaa2, Acadm, Acox1, Bdh1, Ehhadh, Hmgcs2, Ppara*) measured in liver tissue at 10 days post tumor implantation in *ad libitum* fed mice. n = 3 female, 3 male (sham/WT); 3 female, 2 male (sham/IL-6^−/-^); 3 female, 3 male (PDAC/WT); 4 female, 2 male (PDAC/IL-6^−/-^). Error bars represent SEM. All analyses for 2×2 studies were statistically tested with a full effects model 2-way ANOVA with Tukey correction for multiple comparisons. Octanoate challenge was statistically tested with repeated measures ANOVA with Šidák correction for multiple comparisons. Pairwise comparisons tested with parametric, unpaired t-test. **** p<0.0001, ***p<0.001, **p<0.01, *p<0.05. Independent biological replicates are indicated by individual points on bar graphs, unless noted otherwise.

We then evaluated the effect of IL-6 signaling on ketogenesis by subjecting WT and IL-6^−/-^ PDAC mice to an octanoate challenge after a 16 h fast (Figure 4L). First, we found that IL-6^−/-^ PDAC mice had higher fasting ketones than WT PDAC mice. Furthermore, IL-6^−/-^ mice were able to maintain blood ketones at a higher level than WT mice post octanoate challenge (Figure 4M-N). This was further supported by qPCR analysis of hepatic lipid metabolism enzymes that showed restored expression of *Acadm, Acadl, Acox1*, and *Acaa2* in IL-6^−/-^ mice, as compared to WT PDAC mice (Figure 4O). These data indicate that alterations in hepatic lipid metabolism are dependent upon IL-6 signaling, identifying a new site of action for IL-6 in cachexia.

In cachexia, IL-6 is often thought of as a tumor-derived factor, although both IL-6 and a soluble form of its receptor are produced by other tissues and involved in cancer cachexia (Rupert *et al*., 2021). To identify the source of IL-6, we screened tissues in PDAC mice and found that pancreas (tumor) had a significant increase in IL-6 protein in PDAC mice compared to sham (Figure 4D, S4A-B). The absence of IL-6 protein from the pancreas/tumor lysate from IL-6^−/-^ mice supports our conclusion that our KPC cells do not produce IL-6 (Figure 4C-D). This led us to further investigate the intra-tumoral cell population producing IL-6. We identified podoplanin+ (PDPN) cancer-associated fibroblasts, and CD45+ immune cells had the highest levels of IL-6, and that CD45+ immune cells were the most abundant cell population quantified (Figure S4C-D). Within the CD45+ population, CD11b+ (myeloid) cells had the highest IL-6 signal. A mixed population of MHCII negative and positive cells are the major producers of IL-6 in the PDAC microenvironment (Figure S4E-F).

### Hepatic STAT3 signaling mediates tumor-derived IL-6 signals, liver metabolism, and PDAC cachexia

Because hepatocytes express IL-6 receptor (IL-6R) and IL-6 is known to act on the liver via activation of the STAT3 signaling pathway (Heinrich et al., 1998; Lee et al., 2019), we hypothesized that IL-6 may impact hepatic metabolism by directly acting on hepatocytes. To test this, we generated hepatocyte-specific STAT3 knock out (Li-*Stat3*^−/-^) mice and tested the effect of PDAC on liver metabolism and cachexia generation. We confirmed that STAT3 expression was ablated in all Li-*Stat3*^−/-^ mice. Using ROSAmT/mG;Albumin-Cre reporter mice, we validated that expression of cre recombinase driven by the Alb-cre was restricted to hepatocytes (Figure S5A-E). We first challenged Li-*Stat3*^−/-^ mice and littermate controls with PDAC or sham injection, and pair fed sham animals within each genotype to eliminate effects of food intake (Figure 5A). Both *Socs3* expression and STAT3 phosphorylation were increased in PDAC-bearing littermate control mice, but not Li-*Stat3*^−/-^ mice, verifying that STAT3 activity was ablated in the livers of Li-*Stat3*^−/-^ mice (Figure 5B, S5F-G). Circulating IL-6 was elevated to the same degree in both PDAC groups indicating that the effects we see are specifically due to changes in liver signaling, not changes in circulating IL-6 (Figure 5C). We did not see any significant differences in cumulative food intake, daily body mass, or tumor, gWAT and liver masses between genotypes or PDAC condition (Figure 5D-F, S5H-I). Notably, hepatocyte-specific STAT3 deletion completely reversed PDAC-associated gastrocnemius muscle wasting and prevented upregulation of muscle catabolism genes (*Fbxo32, Foxo1, Trim63*) (Figure 5G-H). Furthermore, as in the IL-6^−/-^ mice, we were interested to see if the reversal in cachexia development was tied to changes in hepatic lipid metabolism. We found that Li-*Stat3*^−/-^ PDAC mice had restored expression of several genes related to beta oxidation and ketogenesis (*Acadm, Acadl, Bdh1*, and *Ehhadh*) (Figure 5I). Finally, we compared the ketogenic potential of Li-*Stat3*^−/-^ PDAC to WT PDAC mice by subjecting them to an octanoate challenge (Figure 5J). Li-*Stat3*^−/-^ PDAC mice had statistically higher baseline fasting ketone levels, and significantly higher sustained blood ketone levels after octanoate challenge, relative to WT PDAC mice (Figure 5K-L). These data suggest that the liver is a primary site of action for IL-6 in the development of PDAC cachexia.

**Figure 5.**
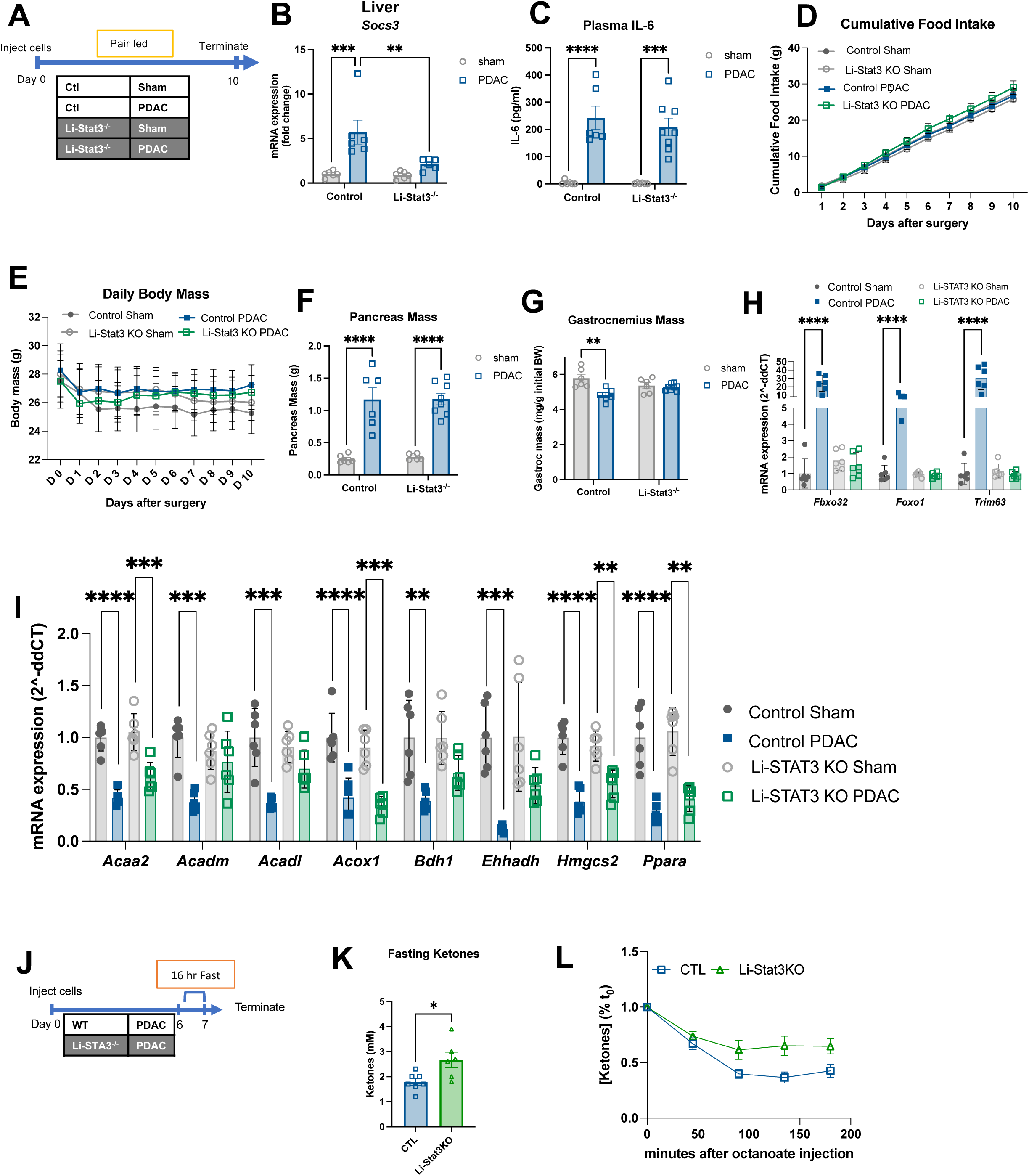
Hepatic STAT3 mediates IL-6 signaling, liver metabolism, and PDAC cachexia. (A) Schema for panels B-I. n = 3 female, 3 male (sham/control); 2 female, 4 male (sham/ Li-*Stat3*^−/-^); 2 female, 4 male (PDAC/control); 3 female, 5 male (PDAC/ Li-*Stat3*^−/-^). (B) qPCR analysis of *Socs3* expression in whole liver of WT and Li-*Stat3*^−/-^ mice. (C) Plasma IL-6 concentration at termination. (D) Cumulative food intake over 10 days of study. (E) Body mass over course of study. (F) Pancreas mass at termination of PDAC and sham animals. (G) Gastrocnemius muscle mass at termination, normalized to initial body weight. (H) qPCR analysis of muscle atrophy genes (*Foxo1, Fbxo32, Trim63*) measured in gastrocnemius muscle. (I) qPCR analysis of lipid metabolism genes (*Acaa2, Acadm, Acox1, Bdh1, Ehhadh, Hmgcs2, Ppara*) measured in liver tissue. (J) Schema showing the 16-hour fast paradigm used in panels K-L. n = 3 female, 3 male (PDAC/Li-*Stat3*^−/-^); 4 female, 3 male (PDAC/control). (K) Fasting ketones in PDAC mice as baseline for octanoate challenge. (L) Blood ketone levels in PDAC mice in response to octanoate challenge after 16 h fast. Time x genotype interaction F_(4, 44)_ = 4.09, p = 0.007. Error bars represent SEM. All analyses for 2×2 studies were statistically tested with a full effects model 2-way ANOVA with Tukey correction for multiple comparisons. Octanoate challenge was statistically tested with repeated measures ANOVA with Šidák correction for multiple comparisons. Pairwise comparisons statistically tested with parametric, unpaired t-test. **** p<0.0001, ***p<0.001, **p<0.01, *p<0.05. Independent biological replicates are indicated by individual points on bar graphs, unless noted otherwise.

## DISCUSSION

We sought to understand the biology underlying the initial tissue wasting in early PDAC cachexia. Our data show that: (1) early-stage cachexia is associated with increased sensitivity to nutritional stress and suppressed lipid oxidation and ketogenesis in the liver, (2) hepatic STAT3 activation by tumor-derived IL-6 signaling is a likely mechanism for driving cachexia progression, and (3) ketogenic diet is sufficient to prevent muscle loss in PDAC mice and ketone body supplementation acts directly on myotubes to prevent atrophy *in vitro*.

Although the etiology of weight loss in PDAC is multifactorial, we identified an early cachexia phenotype, which is defined by the specific vulnerability of PDAC mice to caloric deficiency. Moderate nutrient deprivation was sufficient to induce muscle wasting in mice before overt symptoms of cachexia (muscle loss, weight loss, or anorexia) are present, reflecting one of the key differences between cachexia and pure undernutrition. Interestingly, sensitivity to nutritional deficit was not due to hypermetabolism, or impaired nutrient absorption. Instead, the data pointed toward dysfunction in the adaptive hepatic metabolic response, implicating the liver as a central mediator of cachexia (Lieffers *et al*., 2009; Thibaut et al., 2021). Humans and mice are consistently cycling through fasting and feeding states over the course of the day and extended periods of fasting are especially apparent during sleep hours (Gill and Panda, 2015). For cachectic patients, this presents a daily nutritional stress that can drive progressive lean wasting. The canonical metabolic response to fasting is to utilize stored lipids, and, as a result, humans experience the highest levels of blood ketones during their sleep hours (Masi et al., 2022; McGarry and Foster, 1971). In the absence of lipid oxidation, the liver must utilize other substrates for gluconeogenic and ketogenic metabolism, such as glycogen and amino acids mobilized from the muscle (Felig, 1973; Rémésy and Demigné, 1983). Many have speculated that a reliance on muscle breakdown to supply energy to the body is one contributor to cachexia progression, and our data provide one mechanism by which such a reliance could develop (Argilés and Azcón-Bieto, 1988; Leij-Halfwerk et al., 2000).

Notably, STAT3 deletion from hepatocytes was sufficient to reverse muscle catabolism and restore ketogenesis even in the presence of circulating IL-6. Although previous work has linked both lipid mobilization and liver function to the maintenance of skeletal muscle mass during undernutrition, this provides the first direct evidence that hepatic lipid metabolism is essential for muscle preservation in cachexia. STAT3 is the primary effector of IL-6 signaling, and cachexia in this model is IL-6-dependent. Our combined knockout approaches reveal the liver to be a novel target for IL-6 signaling in cachexia. To our knowledge, this is also the first evidence that hepatic STAT3 activation could suppress lipid oxidation. Existing literature shows that constitutive hepatic STAT3 activation suppresses the gluconeogenic genes *Pck1* and *G6pc*, which are similarly upregulated in a low nutrient context (Inoue *et al*., 2004). Hepatic STAT3 activation via leptin signaling also inhibits *Srebp-1* (a control point for fatty acid synthesis), which can prevent steatosis in obesity models (Gurzov et al., 2016). Despite the apparent deficit in beta oxidation, PDAC mice did not exhibit fatty liver, which may be explained by this anti-lipogenic effect of STAT3 activation. This provides evidence that STAT3 can transcriptionally repress lipid metabolism in the liver and that repressive control of hepatic adaptive metabolism is conserved across metabolic syndromes.

Impaired ketogenesis associates with early-stage cachexia and the data presented here indicate that increased ketone levels provide a direct benefit to muscle tissue, as evidenced by retained muscle mass and decreased myotube atrophy. Although various methods of ketogenic supplementation have been investigated as anti-cancer and anti-cachectic interventions, the field still lacks a concrete understanding of the benefits, or lack-there-of, for this intervention (Cortez and Mackenzie, 2021). Other recent works support the observation that ketogenic diet, or restored ketone production via fenofibrate treatment are sufficient to prevent muscle loss, with varying effects on the tumor (Goncalves *et al*., 2018; Koutnik et al., 2020; Shukla *et al*., 2014). Notably, Shukla *et al* show that media from BHB-treated cancer cells induces a lower level of atrophy than media from untreated cancer cells (Shukla *et al*., 2014). Furthermore, Thomsen *et al* present human data showing that infusion with BHB reduces muscle protein catabolism, which supports our *in vitro* observations (Thomsen et al., 2018). However, a separate, recent study in NSCLC showed that ketogenic diet was not sufficient to prevent tumor progression and cachexia development (Langer *et al*., 2022). These apparently contradictory responses to ketogenic diet highlight the likelihood that cachexia can result from different underlying mechanisms, which may be susceptible to unique therapeutic approaches. Indeed, not all patients with cachexia exhibit elevated levels of IL-6, and the levels of other cachexia mediators, such as growth and differentiation factor-15 or tumor necrosis factor, are not uniformly elevated across all patients with cachexia (Bauskin et al., 2006; Johnen et al., 2007; Okada *et al*., 1998; Palmquist et al., 2020; Ramsey *et al*., 2019; Suh *et al*., 2013). Furthermore, the window to see a benefit in skeletal muscle from ketogenic diet may be limited, as our study only evaluated mice at an early phase of cachexia. Particularly in survival studies, if ketogenic diet does not provide benefit to survival, it may be difficult to observe benefit in muscle retention at animals’ humane endpoints. Furthermore, a diverse array of inflammatory signals is known to contribute to cachexia, and it is possible that different inflammatory states impact the efficacy of ketogenic diet as a cachexia intervention. Clinically, it is important to consider that retained muscle mass, even for a limited time early in cancer diagnosis, could provide substantial benefits to patients by improving quality of life and increasing tolerability of anti-tumor interventions.

There are several notable limitations of this study. First, we utilize an OT-PDAC model with a relatively short experimental timeline. Although we are using this model to provide evidence for metabolic changes early in the development of cachexia, cachectic patients experience early-stage cachexia for much longer than the 7-10 days represented in our studies. The limited timeline in our studies also limits the length of exposure to ketogenic diet intervention. Thus, we cannot provide evidence for the efficacy of ketogenic diet for longer timeframes in cachexia treatment. The effects of ketogenic diet, IL-6 knock out, and hepatocyte-specific STAT3 knock out in cachexia development warrant further investigation in other models of PDAC cachexia. Our work with IL-6 and hepatocyte-specific STAT3 knock out mice provides convincing evidence that IL-6 signaling via STAT3 is the primary driver of hepatic defects. However, other activators of STAT3, such as leukemia inhibitory factor (LIF), leptin, and oncostatin M (OSM) may also meaningfully activate STAT3 in PDAC (Wrona et al., 2021; Zimmers et al., 2016). We also did not consider how IL-6 receptor (IL-6R) and IL-6 trans signaling might contribute to cachexia development in our model, which would be an important future consideration as the liver is a source of IL-6R and trans-signaling is thought to mediate the pro-inflammatory effects of IL-6 (Rupert *et al*., 2021; Schaper and Rose-John, 2015). Lastly, our *in vitro* data show that ketones elicit anti-atrophy effects on cultured myotubes. We do not see reductions in key muscle catabolism genes associated with the observed rescue in myotube size. Thus, the relative effects of ketones on skeletal muscle catabolic and anabolic processes and the contribution of each to skeletal muscle preservation is not known.

Early PDAC cachexia is defined by an increased susceptibility to nutritional stress and dysfunctional response to energy loss. Central to this is the failure of the liver to increase ketogenesis in PDAC mice. We observed global downregulation of lipid oxidation genes in the livers of PDAC mice, which was reversed by three interventions: ketogenic diet, whole body IL-6 knock out, and hepatocyte-specific STAT3 knock out. Our work defines a novel role for hepatic metabolic reprogramming in PDAC cachexia and identifies a new mechanism by which IL-6 may influence wasting. Ultimately, this work highlights the importance of understanding nutritional and metabolic states in cachectic patients, and points toward ketogenic diet as a potential adjuvant nutritional intervention for cachectic patients.

## ACKNOWLEDGEMENTS

We thank all members of the Aaron Grossberg and Daniel Marks labs for their helpful discussion and suggestions. Graphical schematics were made using BioRender (BioRender.com). Liver tissue was embedded and sectioned by the Histopathology Shared Resource for pathology studies, which is supported in part by the University Shared Resource Program at Oregon Health and Sciences University and the Knight Cancer Institute (P30 CA069533 and P30 CA069533 13S5). Flow cytometry experiments were performed in the OHSU Flow Cytometry Shared Resource, this shared resource is supported, in part, by the OHSU Knight Cancer Institute NCI Cancer Center Support Grant P30CA069533. SMK was supported by the Brenden-Colson Center Pancreatic Disease Training Fellowship. RE receives funding from the National Cancer Institute (CA256179); the American Association for Cancer Research – MPM Oncology Cancer Research Grant; Conquer Cancer, the American Society of Clinical Oncology Foundation; and Pancreatic Cancer Action Network. AJG receives funding from the National Cancer Institute (CA245188 and CA264133), the Brenden-Colson Center for Pancreatic Care and the Oregon Pancreas Tissue Registry, and the Knight Cancer Institute.

## AUTHOR CONTRIBUTIONS

Conceptualization, PCAW, KP, AJG. Methodology, PCAW, HM, KP, JD, AQB, RE, SMK, AJG. Validation, PCAW, HM, KP, JD. Formal Analysis, PCAW, HM, KP, JD, AQB, BLW, SMK, AJG. Investigation, PCAW, HM, KP, JD, AQB, BLW, AJG. Writing—Original Draft, PCAW. Writing – Review and Editing, PCAW, HM, AJG. Visualization, PCAW, AJG. Supervision, RE, AJG. Project Administration, RE, AJG. Funding Acquisition, AJG. All authors approved this manuscript.

## INCLUSION AND DIVERSITY STATEMENT

We support inclusive, diverse, and equitable conduct of research.

## STAR METHODS

### RESOURCE AVAILABILITY

#### Lead contact

Further information and requests for resources and reagents should be directed to and will be fulfilled by the lead contact, Aaron J. Grossberg

#### Materials availability

Plasmid sequences and engineered KPC cells generated for the purpose of these studies will be shared upon reasonable request.

#### Data and code availability

- All data reported in this paper will be shared by the lead contact upon request
- This paper does not report original code.
- Any additional information required to reanalyze the data reported in this paper is available from the lead contact upon request.

## EXPERIMENTAL MODELS AND SUBJECT DETAILS

### Animal Studies

#### Husbandry

C57BL/6J (WT, JAX 000664), B6.129S2-*Il6*^*tm1Kopf*^/J (IL-6^−/-^, JAX 002650), B6N.Cg-*Speer6-ps1* ^*Tg(Alb-cre)21Mgn*^ /J (Alb-Cre, JAX 018961), B6.129S1-*Stat3* ^*tm1Xyfu*^ /J (STAT3 fl/fl, JAX 016923), STOCK *Gt(ROSA)26Sor* ^*tm4(ACTB-tdTomato,-EGFP)Luo*^ /J (ROSA^mT/mG^, JAX 007576) mice were purchased from The Jackson Laboratory (Bar Harbor, ME) and maintained in our animal facility. *Kras* ^*G12D*^ *p53* ^*R172H/+*^*Pdx*-*Cre* (KPC) mice were provided as a gift from Rosalie Sears and have been previously described (Hingorani et al., 2005). All mice were housed and bred in a dedicated mouse room with a temperature 26□°C with a 12-h light/dark cycle and 40% humidity. Animals were provided *ad libitum* access to food and water (5L0D, PicoLab) unless otherwise stated. Mice were genotyped according to the standard protocol from The Jackson Laboratory. Unless otherwise stated, all animals were 12 weeks of age at experiment start and sex is defined in the figure legends. All OT-PDAC study animals were individually housed for acclimation at least 7 days prior to procedure and for the duration of study. All tumor studies followed human endpoints. Mouse studies were conducted in accordance with the National Institutes of Health Guide for the Care and Use of Laboratory animals, and approved by the Institutional Animal Care and Use Committee of Oregon Health & Science University.

#### Orthotopic pancreatic ductal adenocarcinoma (OT-PDAC) implantation

We performed OT-PDAC implantations as previously described (Michaelis *et al*., 2017). A new vial of frozen stock KPC cells was thawed 5 days prior to each implantation. Media change, or cell passaging was performed every day prior to implantation. On the day of implantation, we trypsinized cells, quenched trypsin with serum-containing media, counted cells, and centrifuged cells at 1100 RPM for 6 minutes. We resuspended the cell pellet in a volume of PBS to achieve 1 million cells per 23 uL. We then aliquoted 23 uL per animal into separate tubes and kept the tubes of cell suspension at 37°C until implantation. All animals were anesthetized with isoflurane and placed supine. Abdomen was scrubbed with betadine and a para-midline incision was made in the abdomen to expose pancreas. Cancer cells (1×10^6^ cells in a volume of 30-40 uL)or vehicle (PBS) were injected directly into the pancreatic parenchyma. Pancreas was placed back into position and incision was closed using two sutures (4-0 Polysorb) and two skin staples. Animals recovered on a warming pad before being returned to a clean cage.

#### Feeding schemes

For all animal studies we monitored food intake daily, unless otherwise noted. For OT-PDAC studies where pair-feeding was employed, beginning on day 7, sham mice were pair fed to the mean weight of food consumed the prior day by PDAC animals. For food restriction (FR) studies, daily food allotment was calculated as 50% of their *ad libitum* consumption for one week prior to OT-PDAC implantation. Mice were allowed to each *ad libitum* for 3 days after surgery, then underwent FR for the final 4 days prior to termination. For all fasting interventions, mice were moved into a clean cage and food was withheld for the stated duration. For ketogenic diet (KD) studies, we used KD 93M diet (TD160153.PWD) and nutritionally-matched control 93M diet (CD, TD150345) from Envigo (Indianapolis, IN). Mice were pair-fed to the PDAC group within each diet. KD and CD were stored at 4°C per the manufacturer’s instructions.

#### Euthanasia and tissue collection

All animals were humanely euthanized via cardiac puncture under deep isoflurane anesthesia. After euthanasia, we trans-cardially perfused animals with ice-cold PBS. Then, tissues were dissected and weighed prior to flash freezing in liquid nitrogen, or formalin fixation for histological analysis.

### Human studies

Our study population included 37 patients diagnosed with PDAC who underwent surgical exploration or pancreatectomy at OHSU between March 27, 2012, and June 28, 2018, detailed patient information including sex and age are provided in Table S1. Gender identity of the patients were not collected. This study was approved by the OHSU institutional review board (IRB# 21923), and all patients provided informed consent. Patients were identified from an institutional database of 373 consecutive patients with potentially resect-able PDAC. Skeletal muscle, visceral adipose tissue, and subcutaneous adipose tissue areas were measured from a single axial slice at the L3 lumbar vertebral level of the pre-surgical staging computed tomography scan as previously published (Grossberg et al., 2016; Prado et al., 2008). Scans were manually segmented to include all paraspinal and abdominal wall muscles by a single trained image analyst using Slice-O-Matic software (v.4.3; Tomovision) with verification from a board-certified radiation oncologist. Skeletal muscle cross sectional areas were normalized to the square of height in meters and reported as the skeletal muscle index (SMI). Patients were classified as sarcopenic if SMI < 52.4 cm^2^/m^2^ for men or < 38.5 cm^2^/m^2^ for women based on prior reports suggesting prognostic utility of this dichotomization (Grossberg *et al*., 2016; Prado *et al*., 2008). A convenience sample of sarcopenic and non-sarcopenic patients with PDAC was selected based on the availability of fasting plasma samples collected at the time of surgery. Fasting was defined as patients whose surgeries began prior to 0900 h, as all patients were instructed to take nothing by mouth for at least 8 hours prior to surgery, and compliance was assumed to be highest for surgeries scheduled early in the morning. Control samples were collected from individuals without known malignancy who have either germline genetic mutations conferring elevated risk of developing PDAC or pancreatic cysts. Samples were collected at the time of esophagogastroduodenoscopy as part of pancreatic cancer surveillance. Patients were instructed to take nothing by mouth for 4 h prior to endoscopy, and only samples from procedures scheduled prior to 0900 h were used in analysis.

### Cell lines

#### Growth Conditions and validation

All cells were maintained at 37°C and 5% CO_2_ in a humidified incubator. Media are described for each cell line below. All cell lines were tested in house for mycoplasma using Universal Mycoplasma Detection Kit (30-1012K) and were negative.

#### KPC

Primary pancreatic cancer cell lines were generated from tumors harvested from the *Kras*^*G12D/+*^, *Tp53*^*R172H/+*,^ *Pdx1-Cre* (KPC) mouse, which spontaneously develops PDAC. KPC cells are female and were generously shared by Dr. Elizabeth Jaffee(Foley et al., 2015; Michaelis *et al*., 2017). KPC cells were grown on tissue culture treated dishes in growth media consisting of RPMI (Gibco) with 10% fetal bovine serum (Corning) and 1% penicillin/streptomycin (Gibco).

#### Engineered KPC

KPC cells expressing the surface marker Thy1.1 (CD90.1), blasticidin resistance (BSR), and IL-6 were generated from our stock of KPC cells (female) described above. Platinum-E ecotropic packaging cells were transfected with plasmid DNA encoding MSCV-Ova-T2A-Thy1.1 or MSCV-Thy1.1 as described previously (Eil et al., 2016). Briefly, retroviral supernatants were collected 48 hours following Lipofectamine-mediated transfection. Retroviral supernatants were spun at 2000g for 2 hours at 32°C onto Retronectin (Takara bio) coated 6 well non-tissue culture treated plates. Supernatant was removed and KPC parental tumor cells were plated at 2e6 cells/well. KPC cells were incubated on virus-coated plates for 24 hours and then removed to standard tissue culture flasks containing DMEM media (Gibco) supplemented with 10% fetal bovine serum (Corning). Two days later KPC cells were placed in blasticidin (5ug/mL) containing complete DMEM media to select for transduced cells. Following antibiotic selection, successful transduction was confirmed via flow cytometry staining for Thy1.1 and IL6. KPC-BSR cells were implanted for OT-PDAC as described for parental KPC cells. Continued culturing of KPC-BSR cells was done in selection media described.

#### C2C12

We grew C2C12 cells (ATCC CRL-1772, RRID:CVCL_0188, sex: female) on tissue culture treated dishes in growth media consisting of high glucose DMEM media (Gibco) supplemented with 10% fetal bovine serum (Corning) and 1% penicillin/streptomycin (Gibco). Differentiation media (DM) contains 2% horse serum (Corning) in place of 10% fetal bovine serum. To differentiate C2C12 cells, we changed media to DM, the following day added 50% more DM, and the third day changed media to fresh DM. To complete atrophy assays, on the fourth day, we changed DM to KPC CM (see above), or, as “C2C12 CM control,” left the DM on the cells. For ketone supplementation studies, CM was supplemented with sodium β-hydroxybutyrate (Sigma) at the indicated concentrations.

## METHOD DETAILS

### Blood measurements

Prior to mouse euthanasia, we used commercial handheld keto-and glucometers to measure blood ketones (Keto-Mojo GK+) and glucose (Ascencia Contour Next) on blood collected from tail-nick.

### Brefeldin A injections

For intracellular cytokine staining for flow cytometry, we followed previously published protocols for Golgi transport blockade(Liu and Whitton, 2005). Briefly, each mouse received 0.25 mg Brefeldin A (Selleckchem) injected via tail vein 6 hours prior to tissue collection.

### Cell imaging and quantification

We collected phase-contrast images of C2C12 myotubes at the start and completion of atrophy assays using an AMG EVOS fl cell imaging system. In Fiji (Schindelin et al., 2012), we quantified the degree of myotube atrophy by measuring myotube diameter at 3 points across each myotube, for a total of at least 15 myotubes per treatment. Triplicate measures were averaged to achieve one average diameter per myotube, and these values were averaged to provide a single value per treatment. Experimenters were blinded to condition for image analysis. We collected fluorescent images of stained myotubes using an AxioSkop2+ with AxioCam MRM camera (Zeiss).

### Cell collection for downstream applications

For transcript (qPCR) downstream analysis, we washed C2C12 cells 2 times with PBS, then scraped cells in an adequate volume of PBS to cover the tissue culture plate. We transferred scraped cells into microfuge tubes and spun at 0.7 G x 7 minutes. We removed the supernatant and stored pellets at −80°C.

### Conditioned media collection (KPC CM)

We plated cells at 50,000 cells per cm^2^, and allowed them to adhere overnight. The following day, we washed with PBS (Gibco) and changed the media to C2C12 differentiation media (DMEM, 2% HS, 1%P/S). 24 hours after the media change, we collected the KPC CM, centrifuged it at 1100 RPM for 6 minutes, then sterile filtered through a 0.22 um filter. KPC CM was either used immediately or frozen at −80°C for future use.

### Echo Magnetic Resonance Imaging Body Composition

Lean mass, fat mass, total body water, and free water were measured using whole body magnetic resonance imaging (MRI) prior to tumor injection (day 0) and then on days 3, 7, 10, and 14 after tumor injection (EchoMRI, Houston, TX).

### Fecal Analyses

#### Fecal protein and lipid content

Sham and PDAC mice were individually housed and allowed *ad libitum* access to food for 7 days after orthotopic injection. Mice were placed in clean cages on day 5. On day 7 feces were collected from cage bottom. To assess total fecal protein, 10□mg of feces was resuspended in 500uL lysis buffer (2% SDS, 150□mM NaCl, 0.5□M EDTA), sonicated, and centrifuged at 10,000 rpm for 15 mins at 4ºC. The protein concentration was then assessed using a BCA assay according to the manufacturer’s instructions. To assess total fecal lipid content, 1g feces were pulverized using a pepper grinder, resuspended in 5mL PBS, and lipids were extracted using a modification of the Folch method (Folch et al., 1957), as previously described (Kraus et al., 2015). Briefly, resuspended feces were extracted using choloform:methanol and centrifugation at 1,300 x g for 20 min at RT. Organic phase containing the extracted lipids was collected, and fecal weight was measured after evaporation.

#### Fecal protease activity

Fecal protease activity was measured as previously described (Olson *et al*., 2021). Briefly, fresh feces were collected from the distal colon at the time of sacrifice, 7 days after orthotopic injection. Feces were suspended at 10 mg/mL in Protein Buffer A (0.1% Triton X-100, 0.5-M NaCl, 100-mM CaCl_2_), homogenized and sonicated, then centrifuged at 14,000 RPM for 15□min. 100 uL of supernatant was added to 200 uL of 3% azocasein, and incubated at 37ºC for 1 h. 500 uL of 8% tri-chloroacetic acid was added to each sample, vortexed, centrifuged at 9,000 RPM for 5 min, and the supernatant was assayed for absorbance at 366 nm.

### Flow Cytometry

#### Sample preparation

We collected tumors from mice 10 days post implantation, and tumors were weighed, then placed in PBS on ice. After all tumors were collected, we minced tumor tissue and digested (Helms et al., 2022). After dissociation, we strained tumor suspension through at 100 um filter, and performed ACK lysis.

#### Staining

We stained samples with live/dead stain (1:2000) and surface protein antibodies listed in figure legends (1:200 each), and incubated for 20 minutes room temperature. After staining, we washed samples with FACS buffer and pelleted. For intracellular staining (IL-6), we resuspended cells in antibody diluted 1:200 in FACS buffer and incubated at 4°C overnight. The next day we washed and resuspended cells with FACS buffer prior to analysis.

#### Instrumentation and analysis

All samples were analyzed in the OHSU Flow Cytometry Shared Resource using the Cytek Aurora flow cytometer (Cytek Biosystems), data was analyzed in FlowJo™ v10.8.1.

### Human plasma analytes

Fasting plasma beta-hydroxybutyrate concentrations were measured using a colorimetric assay (Cayman Chemical #700190) per manufacturer’s protocol. Fasting plasma glucose was measured by hand held glucometer (Ascencia Contour Next). Plasma IL-6 was measured by immunoassay (ProQuantum Human IL-6 Immunoassay Kit, ThermoFisher) and run on the ABI 7300 (Applied Biosystems) per manufacturer’s protocol with analyses performed using ProQuantum Software (ThermoFisher).

### Immunostaining

Muscle tissue was prepared by placing in a 30% sucrose sink overnight, then cryofreezing in O.C.T. medium (Sakura), and sectioning at 8 um on a cryostat at −18°C. We rehydrated sections in PBS and post-fixed in 4% PFA for 15 minutes. After fixation, all samples were permeabilized in 0.5% TritonX-100, followed by blocking in 3% BSA, 0.2% TritonX-100, and 0.2% Tween-20 in PBS. Primary antibody incubations occurred at RT for 90 min, or 4°C overnight, followed by incubation with secondary antibody at RT for 45 min. All antibodies are listed below. All fluorescent slides were counter stained with DAPI. Coverslips were mounted using ProLong Gold (Molecular Probes) or Fluoroshield (Abcam). We collected images of our stained tissues and cells using a Zeiss AxioSkop and Zeiss Axio Cam mrM.

### Lipid analysis

Blood, liver, and gastrocnemius tissue were collected from PDAC and sham mice fed *ad libitum* and terminated at each of 7 and 10 days after tumor implantation. Mice were either fed *ad libitum* throughout the study or fasted for final 24 hours prior to termination. Blood was collected via percutaneous cardiac puncture and stored on ice in EDTA-treated tubes for 30 min prior to centrifugation at 1,500 x g for 15 minutes. Plasma was collected, snap frozen in liquid nitrogen, and stored at −80ºC for subsequent analysis. Plasma NEFA and TG concentrations were measured using kits according to manufacturer’s protocol (NEFA: Wako Diagnostics 999–34691, 995–34791, 991–34891, and 993–35191; TG: StanBio #2100). For liver and gastrocnemius, lipids were first extracted using a modification of the Folch method (Folch *et al*., 1957). Lipids were extracted in chloroform:methanol (1.4:1), centrifuged at 3,000 RPM for 20 min Organic phase containing the extracted lipids was collected, and crude lipid weight was measured after evaporation in a speedvac. To measure NEFAs and TGs, lipid pellets were resuspended in 10 uL chloroform, then suspended in 100 uL PBS. NEFA and TG concentration were measured using the kits described above, according to manufacturer’s protocol, and final concentration was normalized to tissue input.

### Murine indirect calorimetry

Oxygen consumption (VO_2_), carbon dioxide production (VCO_2_), respiratory exchange ratio (RER), and heat production were measured by indirect calorimetry (Oxymax, Columbus Instruments, Columbus, OH, USA). Mice were housed in separate chambers at 26 ± 1ºC and acclimatized to the chambers for 8 hours a day for 2 days prior to conducting the study. Starting 2 days prior to tumor implantation, mice were placed in the chambers for 27 hours at a time, from 0900 to 1300 h the following day. Recordings were made on days −2, 2, 6, and 9 after tumor implantation. Mice were terminated upon removal from the chamber on day 10. Mice were fed *ad libitum* through the first three days after tumor implantation, then were restricted to 50% of their baseline food intake. Food was placed in cages or chambers at 1700 each day. Samples were recorded every 3 min with room air reference recorded every 30 min and air flow to chambers set at 500 ml/min. Indirect calorimetry measurements reflect periods of movement and inactivity, including both feeding and fasting periods each day.

### Octanoate challenge

Nine days after tumor cell injection, mice in each experimental condition were fasted for 16 h overnight then injected intraperitoneally with sodium octanoate (Sigma), dissolved to 200 mM in 0.9% NaCl, at a volume of 6 ml/kg (Flint *et al*., 2016). Blood ketones and glucose were monitored every 45 minutes at t=0, 45, 90, 135, and 180 min post-injection.

### Oil Red O Staining

Livers from PDAC and sham mice terminated 7 days after orthotopic injection were collected and snap frozen in LN2 after perfusion with ice cold PBS. sections were taken for Oil-Red-O (ORO) staining, as previously described (Mehlem et al., 2013). Briefly, 12 um-thick sections were collected via cryostat and stored at −80°C until staining. After equilibration to room temperature, slides were incubated in 1 mL of ORO working solution at RT for 5 min, then counterstained in Mayer’s hematoxylin for 15 s, before being washed under running water for 30 min, dried, and cover-slipped. Ten bright field images were captured at 20X magnification per sample and quantified in ImageJ using previously described thresholding. Total staining was quantified as integrated density, calculated as the product of *area* and *mean gray value*.

### Plasma analytes

Blood was collected via percutaneous cardiac puncture and stored on ice in EDTA-treated tubes for 30 min prior to centrifugation at 1,500 x g for 15 minutes. Plasma was collected, snap frozen in liquid nitrogen, and stored at −80ºC for subsequent analysis. Plasma concentrations of corticosterone were measured using ELISA (ThermoFisher), per manufacturer specifications, and read on a plate reader (BioTek). Plasma levels of IL-6, insulin, and glucagon were measured using a Milliplex multiplex magnetic bead immunoassay (Millipore) and read on the Luminex 200, per manufacturer instructions. Colorimetric assays were used to measure plasma levels of beta-hydroxybutyrate (Cayman Chemical #700190) and acetoaceteate (Abcam #ab180875).

### Quantitative real-time polymerase chain reaction (qPCR)

We isolated RNA from cell pellets or tissue samples using the E.Z.N.A. Total RNA Kit I (Omega BioTek) at the manufacturer’s specifications, and we prepared cDNA using high-capacity cDNA reverse transcription kit (Applied Biosystems). qPCR was run on the ABI 7300 (Applied Biosystems), using TaqMan Fast Advanced PCR master mix (Applied Biosystems) or SYBR Green master mix (Applied Biosystems). TaqMan primer probes or SYBR primer sequences listed below. Relative expression was calculated using the ΔΔC_t_ method. Statistical analysis was performed on the normally distributed ΔC_t_ values.

### Table of TaqMan qPCR primers

***Table of TaqMan qPCR primers:***
| Gene Name | Supplier | Cat. Number | Used as HK gene in: |
| --- | --- | --- | --- |
| 18S | Thermo | 4333760F | Immortalized cells, muscle, liver |
| *acadm* | IDT | Mm.PT.58.9004416 |  |
| *acox1* | IDT | Mm.PT.58.505.3784 |  |
| *Actb* | Thermo | 4352663 | Adipose |
| *Bdh1* | IDT | Mm.PT.58.17371426 |  |
| *Cpt1* | Thermo | Mm01231183_m1 |  |
| *Ehhadh* | IDT | Mm.PT.58.31548328 |  |
| *Fbxo32* | IDT | Mm.PT.58.7025875 |  |
| *Foxo1* | IDT | Mm.PT.58.6477586 |  |
| *G6pc* | Thermo | Mm00839363_m1 |  |
| *Hmgcs2* | Thermo | Mm00550050_m1 |  |
| *Ppara* | Thermo | Mm00440939_m1 |  |
| *Ppargc1a* | Thermo | Mm01208835_m1 |  |
| *Socs3* | Thermo | Mm00545913_s1 |  |
| *Trim63* | IDT | Mm.PT.58.32840172 |  |
| *Ucp1* | IDT | Mm.PT.58.7088262 |  |

### Table of Sybr qPCR Primers

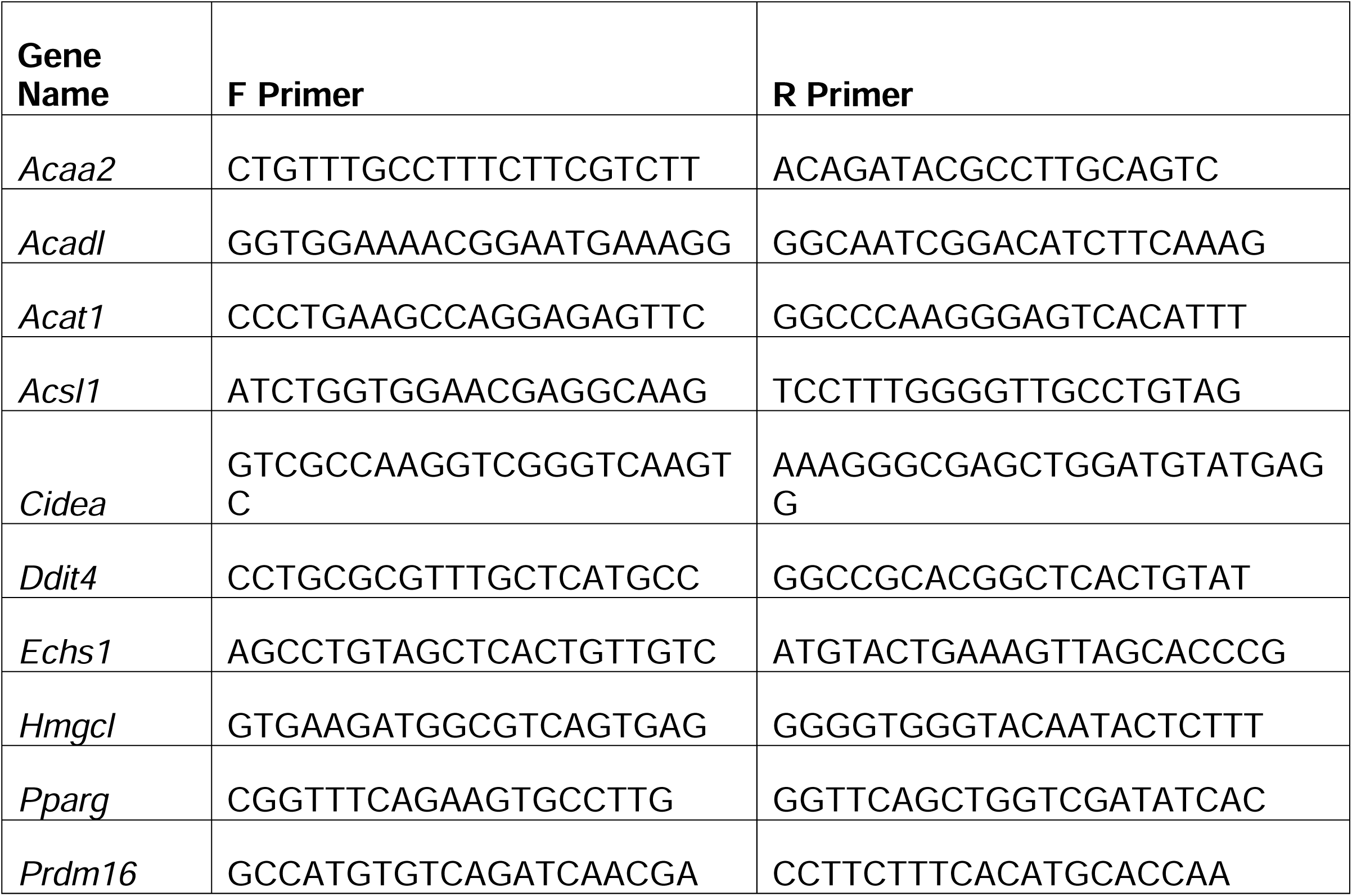

### Western blotting

We extracted protein from snap-frozen tissues by bead homogenization followed by brief sonication. Twenty micrograms of protein were loaded in each lane and run on 10-20% tris glycine gels (Invitrogen). Gels were transferred to PVDF membranes (Millipore) and blocked with 5% BSA for 1□h. Membranes were incubated with primary antibodies overnight at 4□°C with gentle agitation. Blots were then washed with TBST and incubated in secondary antibodies for 1□h prior to imaging (Li-Cor Odyssey imaging system). A complete list of antibodies can be found below.

### Table of antibodies

**Table of antibodies:**
| Protein | Host | Target | Supplier | Cat. Number |
| --- | --- | --- | --- | --- |
| Laminin | Rabbit | Mouse | Sigma | L9393 |
| Myosin Heavy Chain 1E | Mouse | Mouse | DSHB | MF20 |
| STAT3 | Rabbit | Mouse | Cell Signaling | 49045S |
| p-STAT3 | Rabbit | Mouse | Cell Signaling | 9131S |
| IL6 (PE) | Rat | Mouse | BD Bioscience | 562050 |
| CD45.2 (BV786) | Mouse | Mouse | BD Horizon | 563686 |
| PDPN (clone 8.1.1) (FITC) | Syrian Hamster | Mouse | BioLegend | 127415 |
| CD90.1 (clone OX7) (PE-Cy7) | Mouse | Mouse | BD Pharmingen | 561404 |
| CD31 (AF700) | Rat | Mouse | BioLegend | 102443 |
| EpCAM (APC) | Rat | Mouse | Invitrogen | 17-5791-82 |
| CD90.2 (FITC) | Mouse | Mouse | BD Pharmingen | 553004 |
| CD19 (BV480) | Rat | Mouse | BD Bioscience | 566167 |
| CD11b (BV650) | Rat | Mouse | BioLegend | 101239 |
| MHCII (APC Cy7) | Rat | Mouse | BioLegend | 107627 |
| CD4 (PerCP Cy5.5) | Rat | Mouse | BD Pharmingen | 561115 |
| CD8 (AF700) | Rat | Mouse | BD Pharmingen | 557959 |
| Alexa Fluor 488 | Goat | Rabbit | Abcam | ab150077 |
| Alexa Fluor 647 | Goat | Mouse | Abcam | ab150115 |
| DyLight™ 800 4X PEG | Goat | Rabbit | Cell Signaling | 5151P |

## QUANTIFICATION AND STATISTICAL ANALYSIS

Specific statistical tests and sample size for each study is indicated in the figure legends. Error bars in figures show SEM. Comparisons were statistically analyzed using Student’s *t* test, for 2-group comparisons of continuous variables; Pearson χ^2^ for categorical variables; one-way analysis of variance (ANOVA) for 3-group comparisons or two-way ANOVA for factorial studies with multiple comparisons testing using Tukey’s test; or repeated measures ANOVA for within-subjects designs using the Šidák correction for multiple comparisons testing. Myofiber cross-sectional area was compared using nonlinear least squares regression analysis. Statistical analyses were performed using GraphPad Prism (version 9; GraphPad Software Inc) or JMP Pro (version 16; SAS Institute Inc), and graphs were built using GraphPad Prism (GraphPad Software Inc) statistical analysis software. P values are 2 sided with values less than 0.05 regarded as statistically significant.

## SUPPLEMENTAL INFORMATION

## REFERENCES

Acharyya, S., Ladner, K.J., Nelsen, L.L., Damrauer, J., Reiser, P.J., Swoap, S., and Guttridge, D.C. (2004). Cancer cachexia is regulated by selective targeting of skeletal muscle gene products. J Clin Invest 114, 370-378. 10.1172/jci20174.

Argilés, J.M., and Azcón-Bieto, J. (1988). The metabolic environment of cancer. Molecular and cellular biochemistry 81, 3–17.

Arneson-Wissink, P.C., Ducharme, A.M., and Doles, J.D. (2020). A novel transplantable model of lung cancer-associated tissue loss and disrupted muscle regeneration. Skeletal Muscle 10, 1–13.

Babic, A., Schnure, N., Neupane, N.P., Zaman, M.M., Rifai, N., Welch, M.W., Brais, L.K., Rubinson, D.A., Morales-Oyarvide, V., Yuan, C., et al. (2018). Plasma inflammatory cytokines and survival of pancreatic cancer patients. Clin Transl Gastroenterol 9, 145. 10.1038/s41424-018-0008-5.

Baltgalvis, K.A., Berger, F.G., Pena, M.M., Davis, J.M., Muga, S.J., and Carson, J.A. (2008). Interleukin-6 and cachexia in ApcMin/+ mice. Am J Physiol Regul Integr Comp Physiol 294, R393–401. 10.1152/ajpregu.00716.2007.

Baracos, V.E., Martin, L., Korc, M., Guttridge, D.C., and Fearon, K.C. (2018). Cancerassociated cachexia. Nature reviews Disease primers 4, 1–18.

Bauskin, A.R., Brown, D.A., Kuffner, T., Johnen, H., Luo, X.W., Hunter, M., and Breit, S.N. (2006). Role of macrophage inhibitory cytokine-1 in tumorigenesis and diagnosis of cancer. Cancer research 66, 4983–4986.

Bonetto, A., Aydogdu, T., Jin, X., Zhang, Z., Zhan, R., Puzis, L., Koniaris, L.G., and Zimmers, T.A. (2012). JAK/STAT3 pathway inhibition blocks skeletal muscle wasting downstream of IL-6 and in experimental cancer cachexia. American Journal of Physiology-Endocrinology and Metabolism 303, E410–E421.

Bonetto, A., Aydogdu, T., Kunzevitzky, N., Guttridge, D.C., Khuri, S., Koniaris, L.G., and Zimmers, T.A. (2011). STAT3 activation in skeletal muscle links muscle wasting and the acute phase response in cancer cachexia. PLoS One 6, e22538. 10.1371/journal.pone.0022538.

Braun, T.P., Grossberg, A.J., Krasnow, S.M., Levasseur, P.R., Szumowski, M., Zhu, X.X., Maxson, J.E., Knoll, J.G., Barnes, A.P., and Marks, D.L. (2013). Cancer-and endotoxin-induced cachexia require intact glucocorticoid signaling in skeletal muscle. Faseb j 27, 3572–3582. 10.1096/fj.13-230375.

Braun, T.P., Grossberg, A.J., Veleva-Rotse, B.O., Maxson, J.E., Szumowski, M., Barnes, A.P., and Marks, D.L. (2012). Expression of myeloid differentiation factor 88 in neurons is not requisite for the induction of sickness behavior by interleukin-1β. J Neuroinflammation 9, 229. 10.1186/1742-2094-9-229.

Cahill, G.F., Jr. (1970). Starvation in man. N Engl J Med 282, 668–675. 10.1056/nejm197003192821209.

Cortez, N.E., and Mackenzie, G.G. (2021). Ketogenic diets in pancreatic cancer and associated cachexia: Cellular mechanisms and clinical perspectives. Nutrients 13, 3202.

Dalise, S., Tropea, P., Galli, L., Sbrana, A., and Chisari, C. (2020). Muscle function impairment in cancer patients in pre-cachexia stage. European Journal of Translational Myology 30.

Danai, L.V., Babic, A., Rosenthal, M.H., Dennstedt, E.A., Muir, A., Lien, E.C., Mayers, J.R., Tai, K., Lau, A.N., Jones-Sali, P., et al. (2018). Altered exocrine function can drive adipose wasting in early pancreatic cancer. Nature 558, 600–604.

De Benedetti, F., Meazza, C., and Martini, A. (2002). Role of interleukin-6 in growth failure: an animal model. Horm Res 58 Suppl 1, 24–27. 10.1159/000064757.

Ebrahimi, B., Tucker, S.L., Li, D., Abbruzzese, J.L., and Kurzrock, R. (2004). Cytokines in pancreatic carcinoma: correlation with phenotypic characteristics and prognosis. Cancer 101, 2727–2736. 10.1002/cncr.20672.

Eil, R., Vodnala, S.K., Clever, D., Klebanoff, C.A., Sukumar, M., Pan, J.H., Palmer, D.C., Gros, A., Yamamoto, T.N., and Patel, S.J. (2016). Ionic immune suppression within the tumour microenvironment limits T cell effector function. Nature 537, 539–543.

Emmerson, P.J., Wang, F., Du, Y., Liu, Q., Pickard, R.T., Gonciarz, M.D., Coskun, T., Hamang, M.J., Sindelar, D.K., and Ballman, K.K. (2017). The metabolic effects of GDF15 are mediated by the orphan receptor GFRAL. Nature medicine 23, 1215–1219.

Falconer, J.S., Fearon, K., Plester, C.E., Ross, J.A., and Carter, D.C. (1994). Cytokines, the acute-phase response, and resting energy expenditure in cachectic patients with pancreatic cancer. Annals of surgery 219, 325.

Fearon, K., Arends, J., and Baracos, V. (2013). Understanding the mechanisms and treatment options in cancer cachexia. Nature reviews Clinical oncology 10, 90–99.

Fearon, K., Strasser, F., Anker, S.D., Bosaeus, I., Bruera, E., Fainsinger, R.L., Jatoi, A., Loprinzi, C., MacDonald, N., Mantovani, G., et al. (2011). Definition and classification of cancer cachexia: an international consensus. Lancet Oncol 12, 489–495. 10.1016/S1470-2045(10)70218-7.

Fearon, K.C., Glass, D.J., and Guttridge, D.C. (2012). Cancer cachexia: mediators, signaling, and metabolic pathways. Cell metabolism 16, 153–166.

Fearon, K.C., Voss, A.C., Hustead, D.S., and Cancer Cachexia Study, G. (2006). Definition of cancer cachexia: effect of weight loss, reduced food intake, and systemic inflammation on functional status and prognosis. Am J Clin Nutr 83, 1345–1350.

Felig, P. (1973). The glucose-alanine cycle. Metabolism 22, 179–207.

Felig, P., Owen, O.E., Wahren, J., and Cahill, G.F., Jr. (1969). Amino acid metabolism during prolonged starvation. J Clin Invest 48, 584–594. 10.1172/jci106017.

Flatt, J.P., and Quail, J.M. (1981). Effects of liver damage on ketone-body production and nitrogen balance in starved rats. Biochem J 198, 227–230. 10.1042/bj1980227.

Flint, T.R., Janowitz, T., Connell, C.M., Roberts, E.W., Denton, A.E., Coll, A.P., Jodrell, D.I., and Fearon, D.T. (2016). Tumor-induced IL-6 reprograms host metabolism to suppress anti-tumor immunity. Cell metabolism 24, 672–684.

Folch, J., Lees, M., and Sloane Stanley, G.H. (1957). A simple method for the isolation and purification of total lipides from animal tissues. J Biol Chem 226, 497–509.

Foley, K., Rucki, A.A., Xiao, Q., Zhou, D., Leubner, A., Mo, G., Kleponis, J., Wu, A.A., Sharma, R., and Jiang, Q. (2015). Semaphorin 3D autocrine signaling mediates the metastatic role of annexin A2 in pancreatic cancer. Science signaling 8, ra77–ra77.

Fujita, J., Tsujinaka, T., Yano, M., Ebisui, C., Saito, H., Katsume, A., Akamatsu, K., Ohsugi, Y., Shiozaki, H., and Monden, M. (1996). Anti-interleukin-6 receptor antibody prevents muscle atrophy in colon-26 adenocarcinoma-bearing mice with modulation of lysosomal and ATP-ubiquitin-dependent proteolytic pathways. Int J Cancer 68, 637–643. 10.1002/(sici)1097-0215(19961127)68:5<637::Aid-ijc14>3.0.Co;2-z.

Gill, S., and Panda, S. (2015). A smartphone app reveals erratic diurnal eating patterns in humans that can be modulated for health benefits. Cell metabolism 22, 789–798.

Goncalves, M.D., Hwang, S.-K., Pauli, C., Murphy, C.J., Cheng, Z., Hopkins, B.D., Wu, D., Loughran, R.M., Emerling, B.M., and Zhang, G. (2018). Fenofibrate prevents skeletal muscle loss in mice with lung cancer. Proceedings of the National Academy of Sciences 115, E743–E752.

Goodman, M.N., Larsen, P.R., Kaplan, M.M., Aoki, T.T., Young, V.R., and Ruderman, N.B. (1980). Starvation in the rat. II. Effect of age and obesity on protein sparing and fuel metabolism. Am J Physiol 239, E277–e286. 10.1152/ajpendo.1980.239.4.E277.

Goodman, M.N., Lowell, B., Belur, E., and Ruderman, N.B. (1984). Sites of protein conservation and loss during starvation: influence of adiposity. Am J Physiol 246, E383–390. 10.1152/ajpendo.1984.246.5.E383.

Goodman, M.N., and Ruderman, N.B. (1980). Starvation in the rat. I. Effect of age and obesity on organ weights, RNA, DNA, and protein. Am J Physiol 239, E269–e276. 10.1152/ajpendo.1980.239.4.E269.

Grossberg, A.J., Chamchod, S., Fuller, C.D., Mohamed, A.S., Heukelom, J., Eichelberger, H., Kantor, M.E., Hutcheson, K.A., Gunn, G.B., Garden, A.S., et al. (2016). Association of Body Composition With Survival and Locoregional Control of Radiotherapy-Treated Head and Neck Squamous Cell Carcinoma. JAMA Oncol 2, 782–789. 10.1001/jamaoncol.2015.6339.

Grossberg, A.J., Scarlett, J.M., Zhu, X., Bowe, D.D., Batra, A.K., Braun, T.P., and Marks, D.L. (2010). Arcuate nucleus proopiomelanocortin neurons mediate the acute anorectic actions of leukemia inhibitory factor via gp130. Endocrinology 151, 606–616.

Gurzov, E.N., Stanley, W.J., Pappas, E.G., Thomas, H.E., and Gough, D.J. (2016). The JAK/STAT pathway in obesity and diabetes. The FEBS journal 283, 3002–3015.

Heinrich, P.C., Behrmann, I., Müller-Newen, G., Schaper, F., and Graeve, L. (1998). Interleukin-6-type cytokine signalling through the gp130/Jak/STAT pathway. Biochemical journal 334, 297–314.

Helms, E.J., Berry, M.W., Chaw, R.C., DuFort, C.C., Sun, D., Onate, M.K., Oon, C., Bhattacharyya, S., Sanford-Crane, H., and Horton, W. (2022). Mesenchymal lineage heterogeneity underlies nonredundant functions of pancreatic cancer–associated fibroblasts. Cancer discovery 12, 484–501.

Hingorani, S.R., Wang, L., Multani, A.S., Combs, C., Deramaudt, T.B., Hruban, R.H., Rustgi, A.K., Chang, S., and Tuveson, D.A. (2005). Trp53R172H and KrasG12D cooperate to promote chromosomal instability and widely metastatic pancreatic ductal adenocarcinoma in mice. Cancer Cell 7, 469–483. 10.1016/j.ccr.2005.04.023.

Inoue, H., Ogawa, W., Ozaki, M., Haga, S., Matsumoto, M., Furukawa, K., Hashimoto, N., Kido, Y., Mori, T., and Sakaue, H. (2004). Role of STAT-3 in regulation of hepatic gluconeogenic genes and carbohydrate metabolism in vivo. Nature medicine 10, 168–174.

Ji, C., Chen, X., Gao, C., Jiao, L., Wang, J., Xu, G., Fu, H., Guo, X., and Zhao, Y. (2011). IL-6 induces lipolysis and mitochondrial dysfunction, but does not affect insulin-mediated glucose transport in 3T3-L1 adipocytes. J Bioenerg Biomembr 43, 367–375. 10.1007/s10863-011-9361-8.

Johnen, H., Lin, S., Kuffner, T., Brown, D.A., Tsai, V.W.-W., Bauskin, A.R., Wu, L., Pankhurst, G., Jiang, L., and Junankar, S. (2007). Tumor-induced anorexia and weight loss are mediated by the TGF-β superfamily cytokine MIC-1. Nature medicine 13, 1333–1340.

Koutnik, A.P., Poff, A.M., Ward, N.P., DeBlasi, J.M., Soliven, M.A., Romero, M.A., Roberson, P.A., Fox, C.D., Roberts, M.D., and D’Agostino, D.P. (2020). Ketone bodies attenuate wasting in models of atrophy. Journal of cachexia, sarcopenia and muscle 11, 973–996.

Kraus, D., Yang, Q., and Kahn, B.B. (2015). Lipid Extraction from Mouse Feces. Bio Protoc 5. 10.21769/bioprotoc.1375.

Langer, H.T., Ramsamooj, S., Liang, R.J., Grover, R., Hwang, S.-K., and Goncalves, M.D. (2022). Systemic Ketone Replacement Does Not Improve Survival or Cancer Cachexia in Mice With Lung Cancer. Frontiers in Oncology 12.

Lee, J.W., Stone, M.L., Porrett, P.M., Thomas, S.K., Komar, C.A., Li, J.H., Delman, D., Graham, K., Gladney, W.L., and Hua, X. (2019). Hepatocytes direct the formation of a pro-metastatic niche in the liver. Nature 567, 249–252.

Leij-Halfwerk, S., Dagnelie, P.C., van den Berg, J.W.O., Wattimena, J.D.L., Hordijk-Luijk, C.H., and Wilson, J.P. (2000). Weight loss and elevated gluconeogenesis from alanine in lung cancer patients. The American journal of clinical nutrition 71, 583–589.

Lieffers, J.R., Mourtzakis, M., Hall, K.D., McCargar, L.J., Prado, C.M., and Baracos, V.E. (2009). A viscerally driven cachexia syndrome in patients with advanced colorectal cancer: contributions of organ and tumor mass to whole-body energy demands. The American journal of clinical nutrition 89, 1173–1179.

Liu, F., and Whitton, J.L. (2005). Cutting edge: re-evaluating the in vivo cytokine responses of CD8+ T cells during primary and secondary viral infections. The Journal of Immunology 174, 5936–5940.

Llovera, M., García-Martínez, C., López-Soriano, J., Carbó, N., Agell, N., López-Soriano, F.J., and Argiles, J.M. (1998). Role of TNF receptor 1 in protein turnover during cancer cachexia using gene knockout mice. Mol Cell Endocrinol 142, 183–189. 10.1016/s0303-7207(98)00105-1.

Lowell, B.B., and Goodman, M.N. (1987). Protein sparing in skeletal muscle during prolonged starvation. Dependence on lipid fuel availability. Diabetes 36, 14–19. 10.2337/diab.36.1.14.

Martignoni, M.E., Kunze, P., Hildebrandt, W., Künzli, B., Berberat, P., Giese, T., Klöters, O., Hammer, J., Büchler, M.W., Giese, N.A., and Friess, H. (2005). Role of mononuclear cells and inflammatory cytokines in pancreatic cancer-related cachexia. Clin Cancer Res 11, 5802–5808. 10.1158/1078-0432.Ccr-05-0185.

Masi, D., Spoltore, M.E., Rossetti, R., Watanabe, M., Tozzi, R., Caputi, A., Risi, R., Balena, A., Gandini, O., and Mariani, S. (2022). The Influence of Ketone Bodies on Circadian Processes Regarding Appetite, Sleep and Hormone Release: A Systematic Review of the Literature. Nutrients 14, 1410.

McGarry, J.D., and Foster, D.W. (1971). The regulation of ketogenesis from octanoic acid: the role of the tricarboxylic acid cycle and fatty acid synthesis. Journal of Biological Chemistry 246, 1149–1159.

Mehlem, A., Hagberg, C.E., Muhl, L., Eriksson, U., and Falkevall, A. (2013). Imaging of neutral lipids by oil red O for analyzing the metabolic status in health and disease. Nat Protoc 8, 1149–1154. 10.1038/nprot.2013.055.

Michaelis, K.A., Zhu, X., Burfeind, K.G., Krasnow, S.M., Levasseur, P.R., Morgan, T.K., and Marks, D.L. (2017). Establishment and characterization of a novel murine model of pancreatic cancer cachexia. Journal of cachexia, sarcopenia and muscle 8, 824–838.

Mullican, S.E., Lin-Schmidt, X., Chin, C.-N., Chavez, J.A., Furman, J.L., Armstrong, A.A., Beck, S.C., South, V.J., Dinh, T.Q., and Cash-Mason, T.D. (2017). GFRAL is the receptor for GDF15 and the ligand promotes weight loss in mice and nonhuman primates. Nature medicine 23, 1150–1157.

Narsale, A.A., and Carson, J.A. (2014). Role of IL-6 in cachexia–therapeutic implications. Current opinion in supportive and palliative care 8, 321.

Okada, S., Okusaka, T., Ishii, H., Kyogoku, A., Yoshimori, M., Kajimura, N., Yamaguchi, K., and Kakizoe, T. (1998). Elevated serum interleukin-6 levels in patients with pancreatic cancer. Jpn J Clin Oncol 28, 12–15. 10.1093/jjco/28.1.12.

Olson, B., Zhu, X., Norgard, M.A., Levasseur, P.R., Butler, J.T., Buenafe, A., Burfeind, K.G., Michaelis, K.A., Pelz, K.R., and Mendez, H. (2021). Lipocalin 2 mediates appetite suppression during pancreatic cancer cachexia. Nature communications 12, 1–15.

Omlin, A., Blum, D., Wierecky, J., Haile, S.R., Ottery, F.D., and Strasser, F. (2013). Nutrition impact symptoms in advanced cancer patients: frequency and specific interventions, a case–control study. Journal of cachexia, sarcopenia and muscle 4, 55–61.

Onesti, J.K., and Guttridge, D.C. (2014). Inflammation based regulation of cancer cachexia. Biomed Res Int 2014, 168407. 10.1155/2014/168407.

Palmquist, C., Dehlendorff, C., Calatayud, D., Hansen, C.P., Hasselby, J.P., and Johansen, J.S. (2020). Prediction of unresectability and prognosis in patients undergoing surgery on suspicion of pancreatic cancer using carbohydrate antigen 19-9, interleukin 6, and YKL-40. Pancreas 49, 53–61.

Parrilla, R. (1978). Flux of metabolic fuels during starvation in the rat. Pflugers Archiv: European Journal of Physiology 374, 3–7.

Penna, F., Ballarò, R., Martinez-Cristobal, P., Sala, D., Sebastian, D., Busquets, S., Muscaritoli, M., Argilés, J.M., Costelli, P., and Zorzano, A. (2019). Autophagy exacerbates muscle wasting in cancer cachexia and impairs mitochondrial function. Journal of molecular biology 431, 2674–2686.

Pettersen, K., Andersen, S., Degen, S., Tadini, V., Grosjean, J., Hatakeyama, S., Tesfahun, A.N., Moestue, S., Kim, J., Nonstad, U., et al. (2017). Cancer cachexia associates with a systemic autophagy-inducing activity mimicked by cancer cell-derived IL-6 trans-signaling. Sci Rep 7, 2046. 10.1038/s41598-017-02088-2.

Prado, C.M.M., Lieffers, J.R., McCargar, L.J., Reiman, T., Sawyer, M.B., Martin, L., and Baracos, V.E. (2008). Prevalence and clinical implications of sarcopenic obesity in patients with solid tumours of the respiratory and gastrointestinal tracts: a population-based study. The Lancet Oncology 9, 629–635. 10.1016/s1470-2045(08)70153-0.

Puchalska, P., and Crawford, P.A. (2017). Multi-dimensional Roles of Ketone Bodies in Fuel Metabolism, Signaling, and Therapeutics. Cell Metab 25, 262–284. 10.1016/j.cmet.2016.12.022.

Ramsey, M.L., Talbert, E., Ahn, D., Bekaii-Saab, T., Badi, N., Bloomston, P.M., Conwell, D.L., Cruz-Monserrate, Z., Dillhoff, M., Farren, M.R., et al. (2019). Circulating interleukin-6 is associated with disease progression, but not cachexia in pancreatic cancer. Pancreatology 19, 80–87. 10.1016/j.pan.2018.11.002.

Rémésy, C., and Demigné, C. (1983). Changes in availability of glucogenic and ketogenic substrates and liver metabolism in fed or starved rats. Annals of Nutrition and Metabolism 27, 57–70.

Rothman, D.L., Magnusson, I., Katz, L.D., Shulman, R.G., and Shulman, G.I. (1991). Quantitation of hepatic glycogenolysis and gluconeogenesis in fasting humans with 13C NMR. Science 254, 573–576. 10.1126/science.1948033.

Rupert, J.E., Narasimhan, A., Jengelley, D.H., Jiang, Y., Liu, J., Au, E., Silverman, L.M., Sandusky, G., Bonetto, A., Cao, S., et al. (2021). Tumor-derived IL-6 and trans-signaling among tumor, fat, and muscle mediate pancreatic cancer cachexia. Journal of Experimental Medicine 218, e20190450.

Scarlett, J.M., Jobst, E.E., Enriori, P.J., Bowe, D.D., Batra, A.K., Grant, W.F., Cowley, M.A., and Marks, D.L. (2007). Regulation of central melanocortin signaling by interleukin-1β. Endocrinology 148, 4217–4225.

Schaper, F., and Rose-John, S. (2015). Interleukin-6: Biology, signaling and strategies of blockade. Cytokine & growth factor reviews 26, 475–487.

Schindelin, J., Arganda-Carreras, I., Frise, E., Kaynig, V., Longair, M., Pietzsch, T., Preibisch, S., Rueden, C., Saalfeld, S., and Schmid, B. (2012). Fiji: an open-source platform for biological-image analysis. Nature methods 9, 676–682.

Shukla, S.K., Gebregiworgis, T., Purohit, V., Chaika, N.V., Gunda, V., Radhakrishnan, P., Mehla, K., Pipinos, I.I., Powers, R., and Yu, F. (2014). Metabolic reprogramming induced by ketone bodies diminishes pancreatic cancer cachexia. Cancer & metabolism 2, 1–19.

Suh, S.-Y., Choi, Y.S., Yeom, C.H., Kwak, S.M., Yoon, H.M., Kim, D.G., Koh, S.-J., Park, J., Lee, M.A., and Lee, Y.J. (2013). Interleukin-6 but not tumour necrosis factor-alpha predicts survival in patients with advanced cancer. Supportive Care in Cancer 21, 3071–3077.

Talbert, E.E., Cuitino, M.C., Ladner, K.J., Rajasekerea, P.V., Siebert, M., Shakya, R., Leone, G.W., Ostrowski, M.C., Paleo, B., Weisleder, N., et al. (2019). Modeling Human Cancer-induced Cachexia. Cell Rep 28, 1612–1622 e1614. 10.1016/j.celrep.2019.07.016.

Thibaut, M.M., Sboarina, M., Roumain, M., Pötgens, S.A., Neyrinck, A.M., Destrée, F., Gillard, J., Leclercq, I.A., Dachy, G., and Demoulin, J.B. (2021). Inflammation-induced cholestasis in cancer cachexia. Journal of cachexia, sarcopenia and muscle 12, 70–90.

Thomsen, H.H., Rittig, N., Johannsen, M., Møller, A.B., Jørgensen, J.O., Jessen, N., and Møller, N. (2018). Effects of 3-hydroxybutyrate and free fatty acids on muscle protein kinetics and signaling during LPS-induced inflammation in humans: anticatabolic impact of ketone bodies. The American journal of clinical nutrition 108, 857–867.

Trujillo, M.E., Sullivan, S., Harten, I., Schneider, S.H., Greenberg, A.S., and Fried, S.K. (2004). Interleukin-6 regulates human adipose tissue lipid metabolism and leptin production in vitro. J Clin Endocrinol Metab 89, 5577–5582. 10.1210/jc.2004-0603.

von Haehling, S., Anker, M.S., and Anker, S.D. (2016). Prevalence and clinical impact of cachexia in chronic illness in Europe, USA, and Japan: facts and numbers update 2016. Wiley Online Library.

Wrona, E., Potemski, P., Sclafani, F., and Borowiec, M. (2021). Leukemia inhibitory factor: a potential biomarker and therapeutic target in pancreatic cancer. Archivum immunologiae et therapiae experimentalis 69, 1–8.

Xu, P.C., You, M., Yu, S.Y., Luan, Y., Eldani, M., Caffrey, T.C., Grandgenett, P.M., O’Connell, K.A., Shukla, S.K., Kattamuri, C., et al. (2022). Visceral adipose tissue remodeling in pancreatic ductal adenocarcinoma cachexia: the role of activin A signaling. Sci Rep 12, 1659. 10.1038/s41598-022-05660-7.

Zhu, X., Burfeind, K.G., Michaelis, K.A., Braun, T.P., Pelz, K.R., Morgan, T.K., and Marks, D.L. (2018). MyD88 signaling is critical in the development of pancreatic cancer cachexia. Journal of cachexia, sarcopenia & muscle In Press.

Zimmers, T.A., Fishel, M.L., and Bonetto, A. (2016). STAT3 in the systemic inflammation of cancer cachexia. Semin Cell Dev Biol 54, 28–41. 10.1016/j.semcdb.2016.02.009.

Zwickl, H., Hackner, K., Köfeler, H., Krzizek, E.-C., Muqaku, B., Pils, D., Scharnagl, H., Solheim, T.S., Zwickl-Traxler, E., and Pecherstorfer, M. (2020). Reduced LDL-cholesterol and reduced Total cholesterol as potential indicators of early Cancer in male treatment-naive Cancer patients with pre-cachexia and Cachexia. Frontiers in Oncology 10, 1262.

