## Supplemental Figures for "IL-6/STAT3 signaling drives early-stage pancreatic cancer cachexia via suppressed ketogenesis"

Supplemental Figure 1

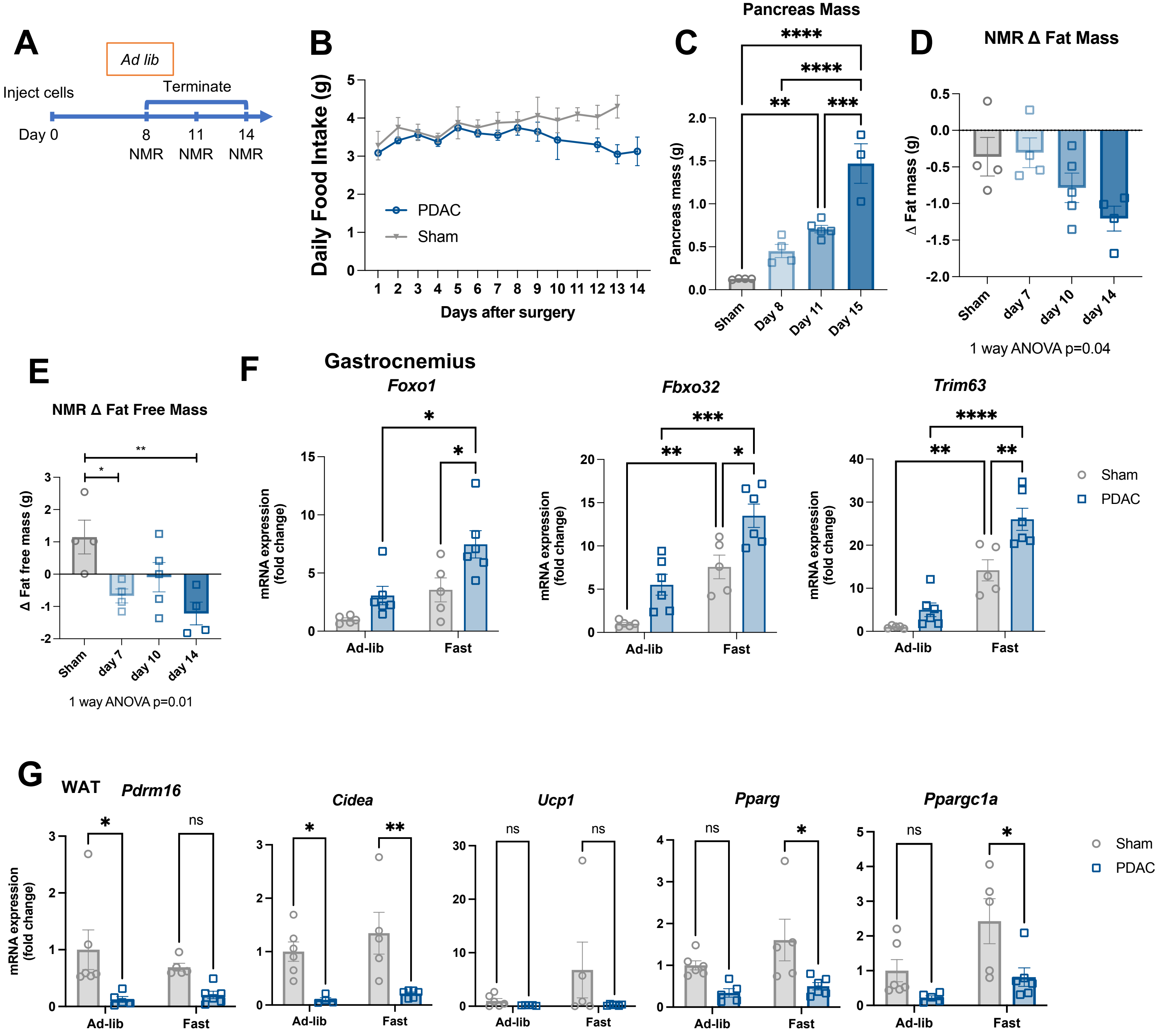

**Supplemental Figure 1. OT-PDAC cachexia progression.** (A) Schematic for panels B-E. n = 3 males (sham); 4 males (PDAC D8); 5 males (PDAC D11); 5 males (PDAC D15). (B)Daily food intake for PDAC and sham animals. (C) Pancreas mass across OT-PDAC progression. (D) EchoMRI measured fat mass relative to baseline across OT-PDAC progression. (E) EchoMRI measured fat-free mass relative to baseline across OT-PDAC progression. (F) qPCR analysis of muscle atrophy genes (*Foxo1*, *Fbxo32*, *Trim63*) measured in gastrocnemius muscle after 16 h fast, 11 d post implantation. n = 5 males (sham groups), 6 males (PDAC groups). (G) qPCR analysis of adipose browning genes (*Cidea*, *Pparg*, *Ppargc1a*, *Prdm16*, *Ucp1*) measured in white adipose tissue, 10 days post implantation after 24 hour fast. n = 6 male mice per group. Error bars represent SEM. All analyses for 2x2 studies were statistically tested with a full effects model 2-way ANOVA and Sidak multiple comparisons test. 4-group analyses tested with one-way ANOVA and Tukey correction for multiple comparisons. **** $p<0.0001$, ***$p<0.001$, **$p<0.01$, *$p<0.05$.

### Supplemental Figure 2

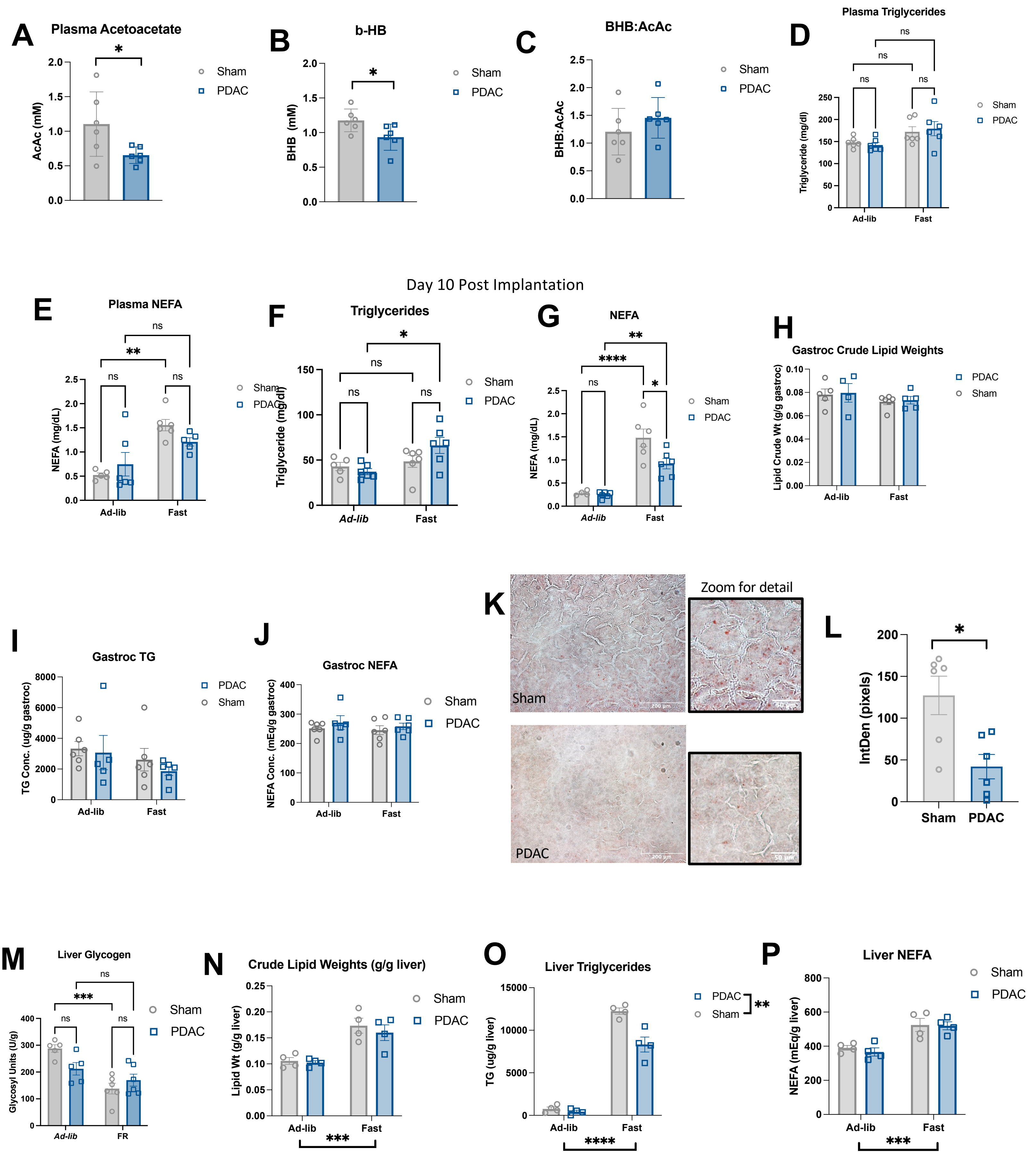

**Supplemental Figure 2. OT-PDAC lowers lipid availability and does not cause lipid accumulation in liver.** (A) Plasma acetoacetate levels (mM) in sham and PDAC mice. (B) Plasma BHB levels (mM) in sham and PDAC mice. (C) Ratio of plasma BHB and acetoacetate levels in sham and PDAC mice. Plasma triglycerides (mg/dL) (A-C, F-G) n = 6 male mice per group. (D) and plasma NEFA (mg/dL) (E) at 7 days post OT-PDAC implantation. Plasma triglycerides (mg/dL) (D-E) 3 female, 3 male mice per group. (F) and plasma NEFA (mg/dL) (G) at 10 days post OT-PDAC implantation. Crude lipid weight (H), triglyceride content (I), and NEFA content (J) in gastrocnemius muscle tissue, normalized to total tissue mass. (H-J) n = 3 female, 3 male (sham groups); 2 female, 3 male (PDAC groups). (K) Oil Red-O staining of liver from sham (top) and PDAC (bottom) with zoom inset to show lipid droplets. (L) Quantification of Oil Red-O staining in liver, measured as integrated density by pixels. n = 3 males, 3 females per group. (M) Liver glycogen content per g of tissue. n = 3 female, 2 male (sham/ad lib); 2 female, 4 male (sham/FR); 3 female, 2 male (PDAC/ad lib); 2 female, 4 male (PDAC/FR). (N) Crude lipid weight, (O) triglyceride content, and (P) NEFA content in liver tissue, normalized to total tissue mass. (N-P) n = 4 male mice per group. Error bars represent SEM. All analyses for 2x2 studies were statistically tested with a full effects model 2-way ANOVA and Sidak multiple comparisons test. Asterisks below the x axis or in the legend indicate significant group effects. Pairwise comparisons statistically tested with parametric, unpaired t-test. **** $p<0.0001$, ***$p<0.001$, **$p<0.01$, *$p<0.05$.

Supplemental Figure 3

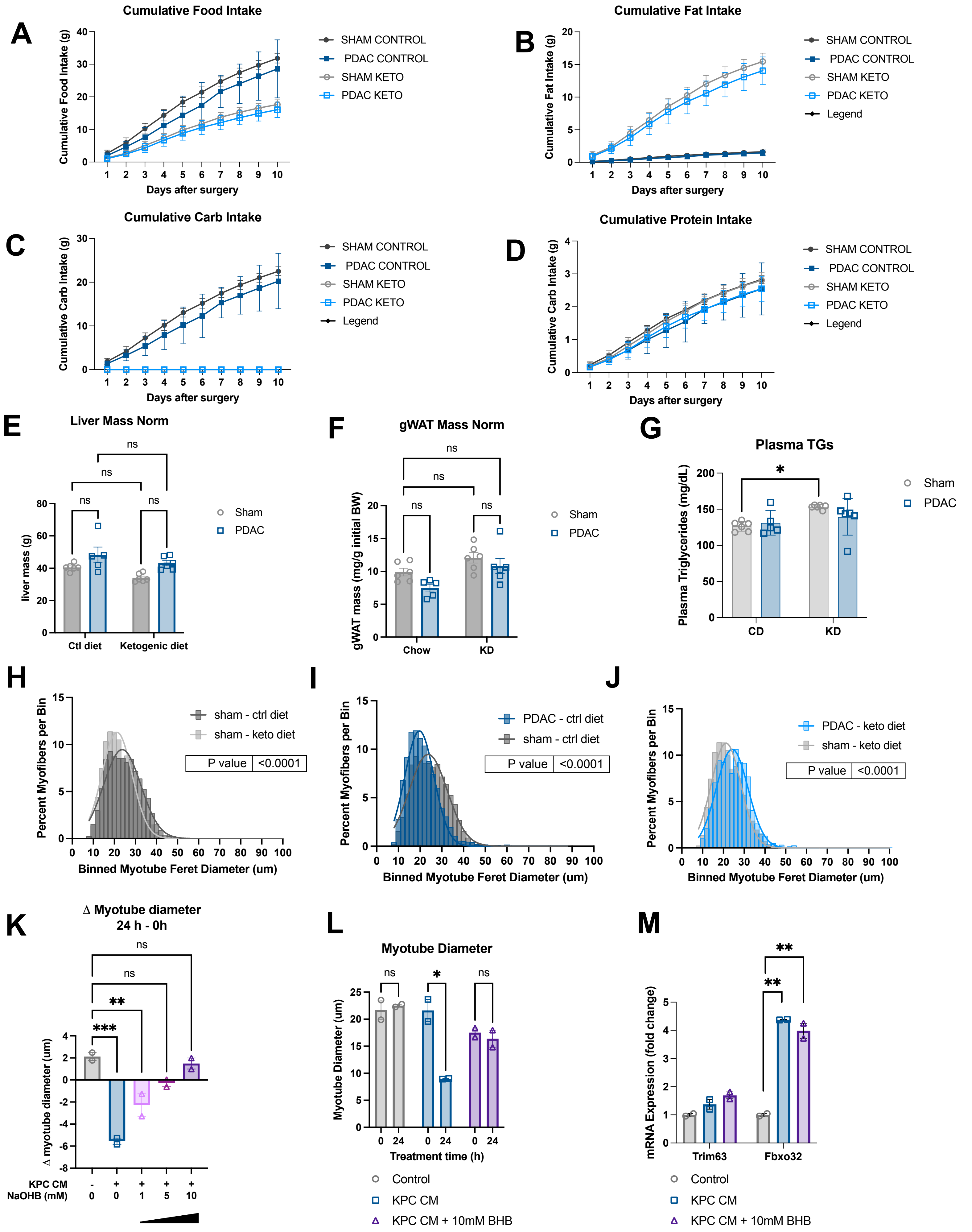

**Supplemental Figure 3. Optimization of *in vitro* atrophy assay and ketogenic diet.** (A-D) Cumulative nutritional intake (g) for mice fed control diet and ketogenic diet**:** food mass (A), fat (B), protein (C), carbohydrate (D). (E) Liver mass at termination. (F) Gonadal white adipose tissue (gWAT) mass at termination normalized to initial body weight. (G) Plasma triglycerides at termination. (H-J) Quantification of minimum feret diameter of myofibers from PDAC and sham mice mice fed control or ketogenic diet. Feret diameters were binned to a histogram and fit with a non-linear regression (Gaussian, least squares regression). P values represented on graph were tested by extra sum-of-squares F test. (A-J) n = 6 male mice (sham/CD, sham/KD, PDAC/KD), 5 male mice (PDAC/CD). (K) Change in C2C12 myotube diameter measured after 24 h exposure to control or KPC cell-conditioned media (KPC CM), supplemented with the indicated concentration of Sodium 3-Hydroxybutyrate (BHB). (L) C2C12 myotube diameter measured at 24 hours post media change to KPC CM +/- 10mM BHB. (M) qPCR analysis of muscle atrophy genes *Trim63* and *Fbxo32* in myotubes after 24 h treatment. (K-M) n = 2 independent wells of 6-well plate per treatment. Error bars represent SEM. All analyses for 2x2 studies were statistically tested with a full effects model 2-way ANOVA and Sidak multiple comparisons test. **** $p<0.0001$, ***$p<0.001$, **$p<0.01$, *$p<0.05$.

| Characteristic | Control | Sarcopenia | | *P*-value |
|---|---|---|---|---|
| | | No | Yes | |
| No. of patients | 14 | 20 | 18 | |
| Mean age at diagosis, years | 62.4 (13.2) | 62.5 (11.4) | 73.6 (9.1) | *0.005* |
| Sex (%) | | | | 0.67 |
| Female | 7 (50) | 13 (65) | 10 (56) | |
| Male | 7 (50) | 7 (35) | 8 (44) | |
| Race (%) | | | | *0.02* |
| Non-white | 4 (29) | 1 (5) | 0 (0) | |
| White | 10 (71) | 19 (95) | 18 (100) | |
| Diagnosis | | | | *<0.0001* |
| Adenocarcinoma | | 20 (100) | 20 (100) | |
| Acute pancreatitis | 3 (21) | | | |
| Chronic pancreatitis | 5 (36) | | | |
| Benign gangioneuroma | 1 (7) | | | |
| Non-malignant mass | 1 (7) | | | |
| Polymorphous lymphocytic population | 1 (7) | | | |
| Serous cystadenoma | 3 (21) | | | |
| Stage (%) | | | | 0.11 |
| Localized | | 3 (15) | 8 (44) | |
| Regional | | 14 (70) | 9 (50) | |
| Metastatic | | 3 (15) | 1 (6) | |
| Grade (%) | | | | 0.80 |
| Well-differentiated | | 1 (5) | 1 (6) | |
| Moderately differentiated | | 10 (50) | 8 (44) | |
| Poorly differentiated | | 4 (20) | 6 (33) | |
| Undetermined | | 5 (25) | 3 (17) | |
| Mean CA19-9 at time of surgery (SD) | | 1036 (2740) | 1154 (3327) | 0.22 |
| Median OS (mo) | Not reached | 18.1 | 17.4 | 0.35 |
| BMI (kg/m$^2$) | 28.2 (5.0) | 30.1 (4.5) | 25.3 (4.8) | *0.01* |
| L3 SMI (cm$^2$/m$^2$) | | 50.8 (1.7) | 37.9 (1.7) | *<0.0001* |
| L3 Muscle CSA (cm$^2$) | | 142.6 (6.6) | 108.8 (6.8) | *0.001* |
| L3 Subcutaneous fat CSA (cm$^2$) | | 259.5 (24.4) | 172.9 (27.5) | *0.02* |
| L3 Visceral fat CSA (cm$^2$) | | 160.6 (19.1) | 121.4 (19.6) | 0.16 |

**Supplemental Table 1. Characteristics of patients with pancreatic cancer.** Continuous variables are reported as mean (SD). Categorical variables reported as number (percentage) at the time of surgery, unless otherwise noted.

| | *Component Percentage by Weight* | |
|---|---|---|
| **Component** | **Ketogenic Diet** | **Control Diet** |
| Carbohydrate | 0% | 70.8% |
| Fat | 67.7% | 5.1% |
| Protein | 15.9% | 8.9% |

**Supplemental Table 2. Description of the macronutrient composition of ketogenic diet versus control diet (percent by weight).**

| Component | g/Kg |
|---|---|
| Casein | 180 |
| DL-Methionine | 2.88 |
| Vegetable Shortening, hydrogenated (Crisco) | 440 |
| Cocoa Butter | 150 |
| Corn Oil | 85 |
| Cellulose | 59.1884 |
| Vitamin Mix, AIN-93-VX w/Cellulose (110068) | 27 |
| Thiamin (81%) | 0.018 |
| Vitamin K1, phylloquinone | 0.0036 |
| Choline Bitartrate | 4.5 |
| Mineral Mix, w/o Ca & P (98057) | 24.1 |
| Calcium Phosphate, dibasic | 17.64 |
| Calcium Carbonate | 9.54 |
| TBHQ, antioxidant | 0.13 |

**Supplemental Table 3. Complete description of components included in ketogenic diet (g component/Kg total food).**

Supplemental Figure 4

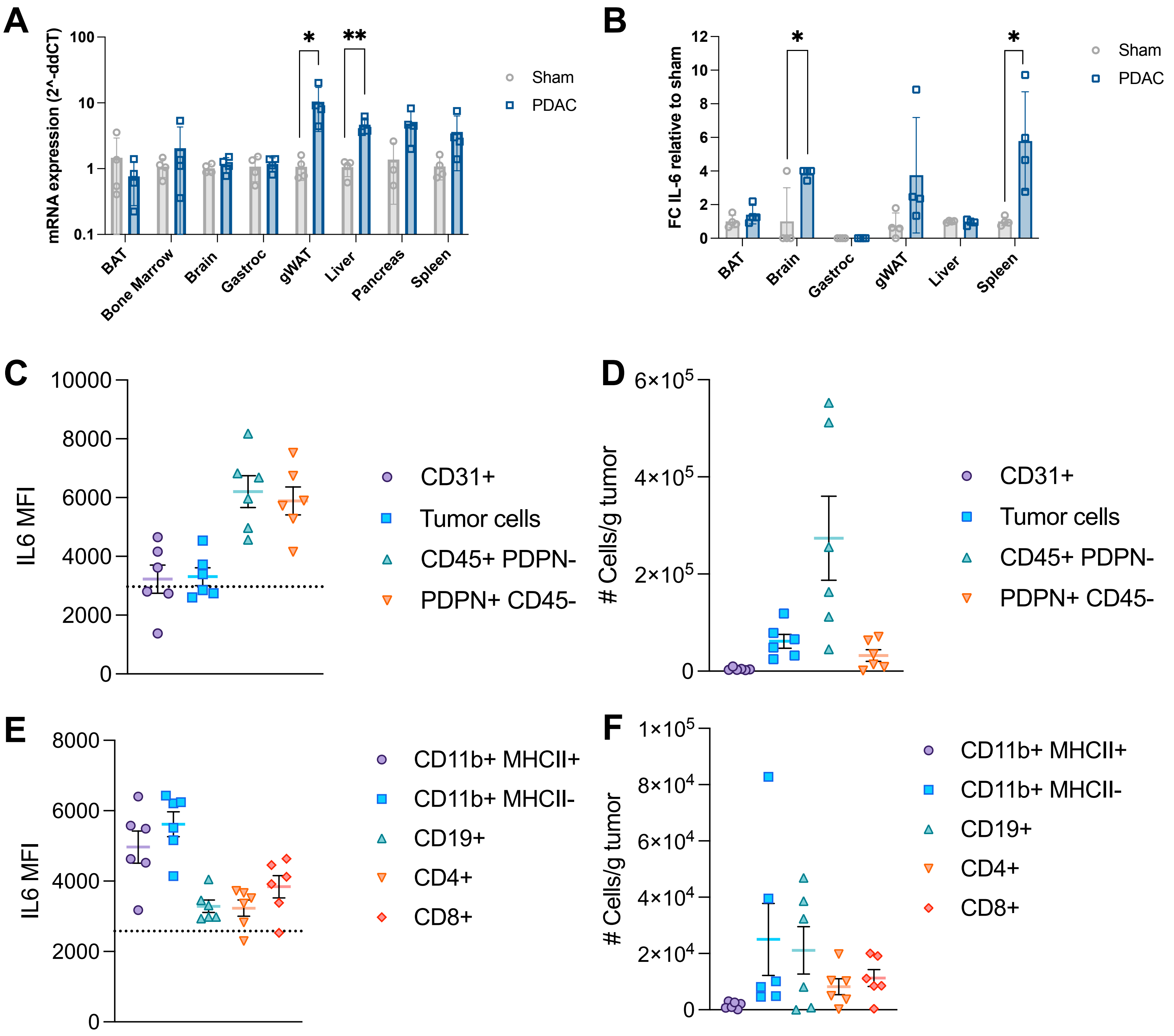

**Supplemental Figure 4. IL-6 is derived from the tumor microenvironment** (A) qPCR for *IL-6* transcript in BAT, bone marrow, brain, gastrocnemius muscle, gWAT, liver, and spleen. (B) IL-6 protein concentration in whole protein lysates from BAT, brain, gastrocnemius muscle (no IL-6 detected), gWAT, liver, and spleen. (A-B) n = 4 male mice per group. (C) Mean fluorescence intensity (MFI) for fluorescently conjugated IL-6 antibody in cell populations isolated from whole pancreas cell suspension. (D) Number of cells per population per gram of tumor, cell number was calculated using CountBright beads. (E) Mean fluorescence intensity (MFI) for fluorescently conjugated IL-6 antibody in CD45+ sub populations isolated from whole pancreas cell suspension. (F) Number of cells per population per gram of tumor, cell number was quantified using CountBright beads. Cell populations are defined as endothelial cells (CD31+), tumor cells (CD90.1+), immune cells (CD45+, PDPN-), and cancer associated fibroblasts (PDPN+ CD45-). Immune cell populations (panels E-F) are within the CD45+ population. (C-F) n = 6 male PDAC mice. Error bars represent SEM. **** $p<0.0001$, ***$p<0.001$, **$p<0.01$, *$p<0.05$.

Supplemental Figure 5

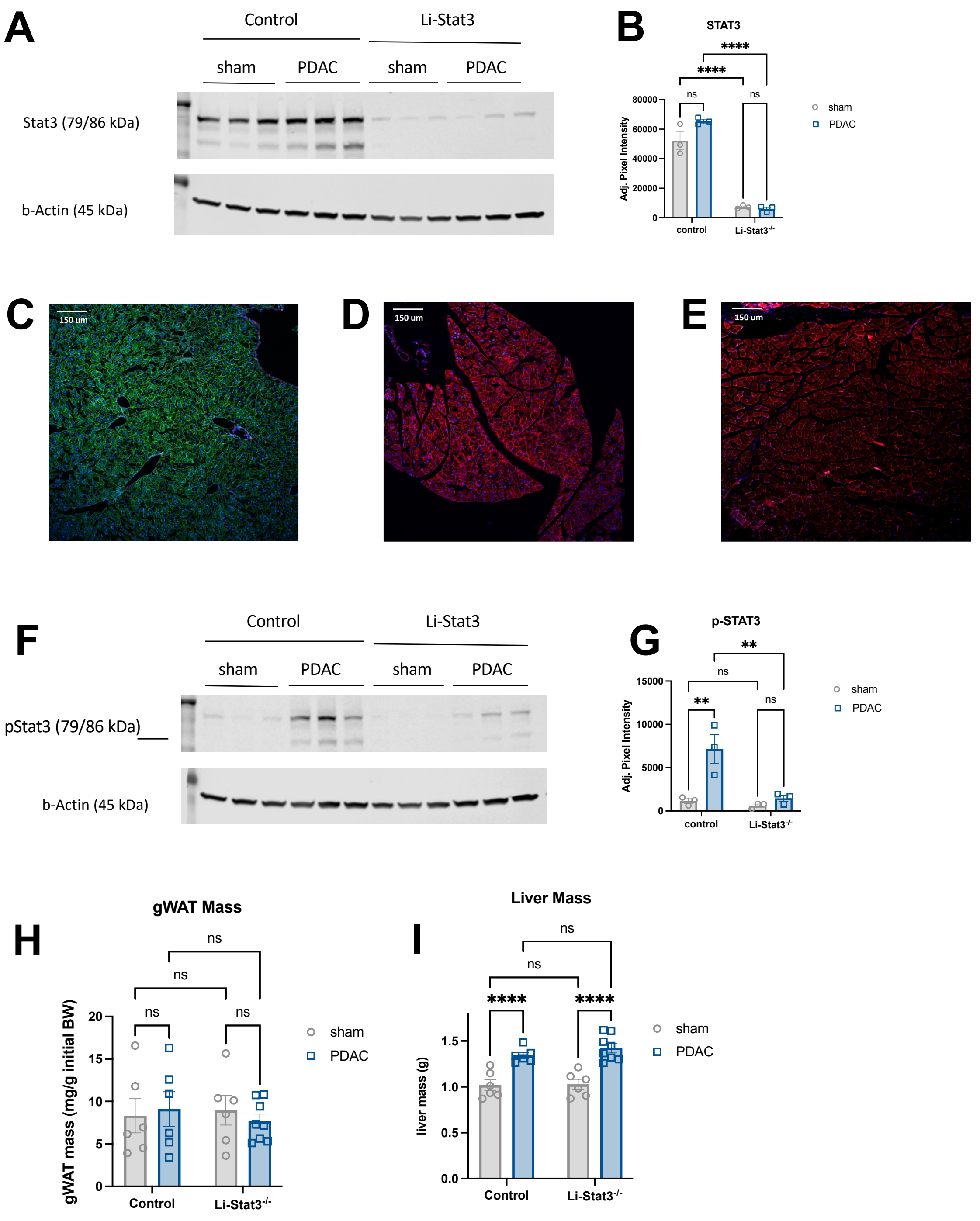

**Supplemental Figure 5. Hepatocyte STAT3 KO is tissue specific.** (A) Western blot for STAT3 (top) and beta-actin (bottom). (B) Densitometry analysis of STAT3 western blot in A, normalized to b-actin expression. (C-E) 3-channel overlay fluorescent images of liver (C), pancreas (D), and gastrocnemius muscle (E) from ROSA$^{mT/}$mG;Albumin-Cre mice, which expresss cell membrane-localized tdTomato in the absence of cre recombinase, and cell membrane-localized EGFP in the presence of cre recombinase. Green is EGFP, red is tdTomato, and blue is DAPI. Scale bar depicts 150 um. (F)Western blot for p-STAT3 (top) and b-actin (bottom). (G) Densitometry analysis of p-STAT3 western blot in C, normalized to b-actin expression. (H) Gonadal white adipose tissue mass at termination, normalized to initial body weight. (I) Liver mass at termination, normalized to initial body weight. (A-B, F-I) n = 3 female, 3 male (sham/control); 2 female, 4 male (sham/ Li-Stat3-/-); 2 female, 4 male (PDAC/control); 3 female, 5 male (PDAC/ Li-Stat3-/-). (C-E) representative images based on n = 3 male mice. Error bars represent SEM. All analyses for 2x2 studies were statistically tested with a full effects model 2-way ANOVA and Sidak multiple comparisons test. **** $p<0.0001$, ***$p<0.001$, **$p<0.01$, *$p<0.05$.
